# Neutrophil terminal programming in the ischemic heart drives fibrosis after myocardial infarction

**DOI:** 10.64898/2026.01.15.699253

**Authors:** Marie Piollet, Giuseppe Rizzo, Ecem T. Sakalli, Jasmin El-Khabbaz, Mathis Gautier, Manuel Gendre, Ludovica Timperi, Anna Rizakou, Sourish Reddy Bandi, Tobias Krammer, Alexander M. Leipold, Jean-Sebastien Silvestre, Antoine-Emmanuel Saliba, Alma Zernecke, Clement Cochain

## Abstract

Following myocardial infarction (MI), the heart undergoes massive neutrophil infiltration characterized by the emergence of distinct subsets, notably a SiglecF^+^ population that accumulates according to specific temporal dynamics. The mechanisms governing cardiac neutrophil heterogeneity and the subsequent functional impact of this diversity on tissue repair following myocardial infarction remain to be elucidated. Using single-cell RNA-sequencing of neutrophils in the heart and peripheral organs of infarcted mice, we here show that while acquisition of the SiglecF^+^ state only fully occurs in the ischemic heart tissue, MI primes neutrophils in the periphery to acquire *Siglecf* and to upregulate receptors for TGFβ and GM-CSF that drive acquisition of the SiglecF^+^ state. Ly6G targeting *in vivo* shifted cardiac neutrophils towards the SiglecF^+^ state at day 3 post-MI, induced the emergence of reprogrammed SiglecF^+^Ly6G^lo^ and *Retnlg*^hi^ neutrophil states at day 5, and was associated with increased fibrosis of the infarct border zone. Mechanistically, Ly6G targeting reshaped the cardiac immune landscape with increased recruitment of pro-fibrotic γδ T cells and monocytes, and SiglecF^+^ neutrophils exerted direct pro-fibrotic effects on fibroblasts in a co-culture system. Altogether, our results indicate that peripheral neutrophil priming combined with their terminal programming towards a SiglecF^+^ state in the ischemic heart drives cardiac fibrosis after MI.

## Introduction

Myocardial infarction (MI) triggers an intense local and systemic inflammatory response playing an essential role in ischemic heart repair and remodeling, and in the development of post-MI heart failure. After MI, the first responders recruited within the heart are neutrophils. Neutrophils initiate tissue repair by clearing dead cells and debris, inducing neovascularization, and producing anti-inflammatory cytokines^1^. They participate in macrophage recruitment within the heart via secretion of cytokines and chemokines^2^, and apoptotic neutrophil phagocytosis drives macrophages towards a pro-resolving phenotype^3,4^. A study using neutrophil depletion in mice induced by anti-Ly6G antibodies further suggested that neutrophils can polarize macrophages towards a reparative phenotype after MI through a Lcn2-dependent paracrine mechanism^5^. On the other hand, neutrophils show detrimental effects by producing reactive oxygen species (ROS) and activating myeloperoxidase^6^, expressing pro-inflammatory cytokines^2^, producing neutrophil extracellular traps (NETs)^7,8^, and by inducing arrhythmias^9^, overall increasing inflammatory signals and leading to pathological myocardial remodeling, as suggested by studies inhibiting recruitment of neutrophils to the heart^6,9–12^.

Recent studies, based on single-cell RNA-seq analyses, identified dynamic accumulation of heterogeneous neutrophil populations within the infarcted heart^13–16^. Notably, our group and others evidenced time-dependent, cardiac tissue-specific appearance of a SiglecF^hi^ neutrophil population^13,16^ with specific functional properties such as increased ROS production. SiglecF^hi^ neutrophils have been described in other contexts such as cancer or kidney failure and associated with enhanced fibrosis^17–19^. Furthermore, we demonstrated that the conventional anti-Ly6G antibody-mediated depletion strategy failed to effectively clear cardiac neutrophils post-MI. Instead, this approach induced a phenotypic shift toward a SiglecF^+^ state, suggesting that previously reported outcomes of neutrophil ‘depletion’ may, in fact, stem from the functional modulation of these cells^16^. However, in the context of MI, the mechanisms underlying the acquisition of the SiglecF^+^ signature within the tissue, and the functional role of SiglecF^+^ neutrophils remain unknown.

Here, we show that after MI, neutrophils are primed in the periphery to acquire *Siglecf* expression and upregulate receptors for tumor growth factor (TGF)β and granulocyte-macrophage colony stimulating factor (GM-CSF) driving acquisition of the SiglecF^+^ state, which fully occurs specifically in the ischemic heart tissue. Shifting the cardiac neutrophil population toward a SiglecF^+^ state using Ly6G antibodies promoted fibrosis through two distinct mechanisms: the direct activation of fibroblasts and the recruitment of pro-fibrotic immune populations, specifically monocytes and γδ T cells.

## Material and methods

### In vivo model of MI and anti-Ly6G targeting

C57BL6/J male and female mice were purchased from Janvier Labs and were enrolled for experiments between 8 and 12 weeks of age. Mice were kept in individually ventilated cages with a 12h light cycle and had free access to food and water. All animal studies and numbers of animals used conform to the Directive 2010/63/EU of the European Parliament and have been approved by the appropriate local authorities (Regierung von Unterfranken, Würzburg, Germany, Akt.-Z. 55.2-DMS-2532-2-865). Myocardial infarction was induced by permanent ligation of the left descendant coronary artery. Briefly, mice received Buprenorphine 0.1mg/kg s.c. as analgesia and were anesthetized with 4% isoflurane inhalation. Once under deep anesthesia, mice were intubated with an endotracheal cannula and placed under mechanical ventilation (Physitemp Instruments inc., TCAT-2LV or VentElite, Harvard apparatus) on a heating pad to maintain body temperature. Mice were maintained under anesthesia with 1.5-2.5% isoflurane during all the procedure. After thoracotomy, the heart was exposed through the 3^rd^ intercostal space. The left descendant coronary artery was visualized and ligated with a 7/0 non resorbable nylon suture (Serapren, CP05341A). The thorax and the skin were closed with 6/0 non resorbable nylon suture thread (Serapren, CP07281A). Mice were injected with buprenorphine 0.1mg/kg s.c. twice daily during 2 days post-surgery. Sham mice underwent the same procedure without the ligation of the left descendant coronary artery. Mice were sacrificed by cervical dislocation under isoflurane anesthesia. For Ly6G targeting, mice received daily intraperitoneal injections of 25µg of rat IgG2aκ anti-mouse Ly6G InVivoPlus™ (BioXCell, BP0075-1) + 50µg of mouse anti-rat IgG2aκ (BioXCell, BE0122) (anti-Ly6G group), and the control group received 25µg of rat IgG2aκ isotype control (BioXCell, BP0089) + 50µg of mouse anti-rat IgG2aκ (BioXCell, BE0122) (isotype group). Injections started 2 days before the induction of MI and lasted until day 7 post-MI.

### Cell preparation for flow cytometry and single cell RNA-seq/CITE-seq analysis

Mice were injected i.v. with 2.5µg anti-CD45.2 clone 104 (various fluorochromes indicated for individual experiments) under isoflurane anesthesia, then sacrificed after 5min by cervical dislocation.

*Heart cells:* The heart was exposed and perfused with PBS+1%FCS, excised, and the infarct, peri-infarct area and adjacent viable myocardium collected, weighed and digested for 45min at 37°C under agitation in RPMI containing 450U/ml collagenase I (Sigma-Aldrich, C0130), 125U/ml Collagenase XI (Sigma-Aldrich, C7657), 60U/ml Hyaluronidase (Sigma-Aldrich, H3506), 60U/ml DNAse (Roche, 11284932001), filtered on a 70µm cell strainer and washed once with PBS+1%FCS at 4°C. Alternatively, the hearts were mechanically dissociated and filtered on a 70µm cell strainer to preserve specific epitopes (CXCR2) for flow cytometry analysis. Absolute cell counts were determined by flow cytometry using Precision Count Beads (Biolegend, 424902). CD45^+^ cell counts were then normalized to the weight of the processed heart sample (CD45^+^ cells/mg of tissue).

*BM cells:* A femur was harvested and bone marrow was isolated by centrifugation and filtered on 70µm cell strainer. Cells were washed once with PBS+1%FCS at 4°C, and ready for further analysis.

*Blood cells:* Before sacrifice, mice were bled under isoflurane-induced deep anesthesia via retro-orbital puncture. The blood was collected in EDTA-covered tube (Starsted, 41.1500.005) and kept at 4°C. Erythrocytes were lysed with lysis buffer (150 mM Ammonium chloride NH4Cl, 10 mM Potassium bicarbonate KHCO3, and 0.1 mM EDTA in distilled water) applied twice for 10min at 4°C and washed with PBS+1%FCS. Remaining blood cells were washed once in PBS+1%FCS at 4°C and ready for further analysis.

*Splenic cells:* the spleen was collected, mechanically dissociated and filtered on a 70µm cell strainer. Erythrocytes were lysed with lysis buffer (150 mM Ammonium chloride NH4Cl, 10 mM Potassium bicarbonate KHCO3, and 0.1 mM EDTA in distilled water) for 10min at 4°C and washed with PBS+1%FCS. Remaining cells were washed once in PBS+1%FCS at 4°C and ready for further analysis.

### Single cell RNA-seq/CITE-seq sample preparation

#### Experiments 1-3: Heart CD45^+^ cells 5 days post-MI after Ly6G targeting

Mice were injected i.v. with 2.5µg anti-CD45.2-BV421 (Biolegend, 109832) under isoflurane anesthesia, then sacrificed after 5min by cervical dislocation. After obtention of a heart cell suspension, the cells were kept at 4°C during all the sample preparation procedure. The cells were washed twice in MACS buffer (PBS+0.5% BSA+2mM EDTA), resuspended in MACS buffer + TruStain FcX™ (Biolegend, 101320) (10µg/ml) and incubated for 10min. The cells were then incubated 15 min with anti-CD45 microbeads (Miltenyi Biotec, 130-052-301) together with anti-CD45.2-AlexaFluor488 (Biolegend, 109815) (Experiments 1 and 2), anti-CD45.2-AlexaFluor700 (Biolegend, 109821) (Experiment 3), then 15min with TotalSeq-A hashtag antibodies for sample multiplexing (one hashtag per experimental animal, see Extended Table 1b for hashtag/sample correspondence for each experiment). Cells were washed twice with MACS buffer, pooled, and CD45^+^ cells were magnetically isolated with LS Columns (Miltenyi Biotec, 130-042-401) and MidiMACS separators (Miltenyi Biotec, 130-042-302). Cells were then washed once, resuspend in PBS+1%BSA, and incubated 20min in a mix containing CITE-seq antibodies (Biolegend, panel in Extended Table 2 CITE-seq TotalSeq-A, see specificities for each experiments) and Fixable Viability Dye e780 (Thermofisher, 65-0865-14, 1:1000). Live/CD45i.v.^−^/CD45^+^ cells were then sorted in PBS+1%BSA with a BD FACS aria III with a 100µm nozzle. Cells were then washed and resuspended in PBS+0.04% ultrapure BSA (ThermoFisher, AM2618), counted and adjusted to a final concentration of 1200 cells/µl, and cells were loaded in 10X Chromium Next GEM Single Cell 3′ Kit (10x Genomics) aiming for 20,000 cells for experiment 1 and 2 and 10x Genomics Chromium Next GEM Single Cell 3′ Kit HT (10x Genomics) aiming for 48,000 cells for experiment 3 and processed according to the manufacturer instructions. Experimental groups and hashtags: Experiment 1: male sham-operated isotype-treated mice (n=3, #1-3); male sham-operated anti-Ly6G-treated mice (n=3, #4-6); male MI-operated isotype-treated mice (n=3, #7-9), male MI-operated anti-Ly6G-treated mice (n=6, #10-15). Experiment 2: female sham-operated isotype-treated mice (n=3, #1-3); female sham-operated anti-Ly6G-treated mice (n=3, #4-6); female MI-operated isotype-treated mice (n=4, #7-10), female MI-operated anti-Ly6G-treated mice (n=5, #11-15). Experiment 3: male MI-operated isotype-treated mice (n=5, #1-5), male MI-operated anti-Ly6G-treated mice (n=5, #6-10).

#### Experiments 4-5: BM, blood, spleen and heart neutrophils 1 day post-MI

Mice were injected i.v. with 2.5µg anti-CD45.2-BV421 (Biolegend, 109832) under isoflurane anesthesia, then sacrificed after 5min by cervical dislocation. After obtention of heart, blood, spleen and bone marrow cells, the cells were kept at 4°C during all the sample preparation procedure. The cells were washed twice in MACS buffer (PBS+0.5% BSA+2mM EDTA), resuspended in MACS buffer + TruStain FcX™ (Biolegend, 101320, 10µg/ml) and incubated for 10min. The heart cells were then incubated 15 min with anti-CD45.2-Alexa488 (Biolegend, 109815) together with anti-CD45 microbeads (Miltenyi Biotec, 130-052-301), then 15min with TotalSeq-B hashtag antibodies for sample multiplexing (one hashtag per experimental animal, see Extended Table 1b for hashtag/sample correspondence for each experiment). Cells were washed twice with MACS buffer, pooled, and CD45^+^ cells were magnetically isolated with LS Columns (Miltenyi Biotec, 130-042-401) and MidiMACS separators (Miltenyi Biotec, 130-042-302). The blood, spleen and bone marrow cells were incubated 15 min with anti-CD45.2-Alexa488 (Biolegend, 109815) then 15min with TotalSeq-B hashtag antibodies for sample multiplexing (one hashtag per experimental sample, see Extended Table 1b). Heart, blood, spleen and bone marrow cells were then washed once, resuspend in PBS+1%BSA, and incubated 20min in a mix containing CITE-seq antibodies (TotalSeq™-B Mouse Universal Cocktail, V1.0, Biolegend, 199902) and Fixable Viability Dye e780 (Thermofisher, 65-0865-14, 1:1000), anti-CD3-PECy7 (Biolegend, 100219, 1:200), anti-NK1.1-PECy7 (Biolegend, 108714, 1:200), CD19-PECy7 (Biolegend, 152417, 1:200) antibodies for flow cytometry cell sorting. Live/CD45i.v.^−^/CD45^+^/CD3^−^NK1.1^−^CD19^−^ heart cells and Live/CD45^+^/CD3^−^NK1.1^−^CD19^−^ spleen, bone marrow and blood cells were then sorted in PBS+1%BSA with a BD FACS aria III with a 100µm nozzle. Cells were then washed and resuspended in PBS+0.04% ultrapure BSA (ThermoFisher, AM2618), counted and adjusted to a final concentration of 1200 cells/µl. Cells were loaded in 10x Genomics Chromium Next GEM Single Cell 3′ HT Kit (10x Genomics) aiming to recover52,000 cells for experiment 4 and 44,000 cells for experiment 5. Experimental groups and hashtags: Experiment 4: male MI-operated mice (n=4, #1-4 for heart and #10-13 for blood cells); male sham-operated mice (n=3, #5-7 for heart and #14-16 for blood cells); male control mice (n=2, #8-9 for heart and #17-18 for blood cells). Experiment 5: male MI-operated mice (n=4, #1-4 for spleen and #10-13 for bone marrow cells); male sham-operated mice (n=3, #5-7 for spleen and #14-16 for bone marrow cells); male control mice (n=2, #8-9 for spleen and #17-18 for bone marrow cells)

#### Experiments 6-9: BM, blood, spleen, heart neutrophils 4 days post-MI

Mice were injected i.v. with 2.5µg anti-CD45.2-APC (Biolegend 109814) under isoflurane anesthesia, then sacrificed after 5min by cervical dislocation. After obtention of heart, blood, spleen and bone marrow cells, the cells were kept at 4°C during all the sample preparation procedure. The cells were washed twice in MACS buffer (PBS+0.5% BSA+2mM EDTA), resuspended in MACS buffer + TruStain FcX™ (Biolegend, 101320, 10µg/ml) and incubated for 10min. The heart cells were then incubated 15 min with anti-CD45.2-Alexa488 (Biolegend, 109815) together with anti-CD45 microbeads (Miltenyi Biotec, 130-052-301), then 15min with TotalSeq-A hashtag antibodies for sample multiplexing (one hashtag per experimental sample, see Extended Table 1b). The cells were washed twice with MACS buffer, pooled, and CD45^+^ cells were magnetically isolated with LS Columns (Miltenyi Biotec, 130-042-401) and MidiMACS separators (Miltenyi Biotec, 130-042-302). Cells were then washed once, resuspend in PBS+1%BSA, and incubated 20min in a mix containing CITE-seq antibodies (Biolegend, panel see Extended Table 2a) and Fixable Viability Dye e780 (Thermofisher, 65-0865-14, 1:1000) for flow cytometry cell sorting. Live/CD45i.v.^−^/CD45^+^ cells were then sorted in PBS1%BSA with a BD FACS aria III with a 100µm nozzle. Cells were then washed and resuspended in PBS+0.04% ultrapure BSA (ThermoFisher, AM2618), counted and adjusted to a final concentration of 1000 cells/µl. Cells were loaded in the 10X Chromium Next GEM Single Cell 3′ HT Kit (10x Genomics) aiming to recover 48,000 cells. The blood, spleen and bone marrow cells were washed twice in MACS buffer (PBS+0.5% BSA+2mM EDTA), resuspended in MACS buffer + TruStain FcX™ (Biolegend, 101320, 10µg/ml) and incubated for 10min. The cells were then incubated 15 min with anti-CD45.2-Alexa488 (Biolegend, 109815) antibody, then 15min with TotalSeq-A hashtag antibodies for sample multiplexing (one hashtag per experimental sample, see Extended Table 1). Cells were washed twice with MACS buffer, pooled, resuspend in PBS+1%BSA, and incubated 20min in a mix containing CITE-seq antibodies (Biolegend, panel see Extended Table 2a) and Fixable Viability Dye e780 (Thermofisher, 65-0865-14, 1:1000), anti-Ly6G-Pacific Blue(Biolegend127612, 1:200) and anti-CD11b-PercpCy5.5 (Biolegend, 101228, 1:200) antibodies for flow cytometry cell sorting. Live/CD45^+^/CD11b^+^/Ly6G^+^, Live/CD45^+^/CD11b^+^/Ly6G^−^and Live/CD45^+^/CD11b^−^ cells were sorted in PBS+1%BSA in the same collection tube at 33% (Live/CD45^+^/CD11b^+^/Ly6G^+^), 33%(Live/CD45^+^/CD11b^+^/Ly6G^−^), 33% (Live/CD45^+^/CD11b^−^) with a BD FACS aria III with a 100µm nozzle. Cells were then washed and resuspended in PBS+0.04% ultrapure BSA (ThermoFisher, AM2618), counted and adjusted to a final concentration of 1000 cells/µl, and cells were loaded in the 10X Chromium Next GEM Single Cell 3′ HT Kit (10x Genomics) aiming for 48,000 cells. Experimental groups and hashtags:Experiment 6 Heart: Male sham-operated mice (n=3, #1-3); female sham-operated mice (n=3, #4-6), male MI-operated mice (n=4, #7-10); female MI-operated mice (n=5, #11-15).Experiment 7 Blood: Male sham-operated mice (n=3, #1-3); female sham-operated mice (n=3, #4-6), male MI-operated mice (n=4, #7-10); female MI-operated mice (n=5, #11-15).Experiment 8 Spleen: Male sham-operated mice (n=3, #1-3); female sham-operated mice (n=3, #4-6), male MI-operated mice (n=4, #7-10); female MI-operated mice (n=5, #11-15). Experiment 9 bone marrow: Male sham-operated mice (n=3, #1-3); female sham-operated mice (n=3, #4-6), male MI-operated mice (n=4, #7-10); female MI-operated mice (n=5, #11-15).

#### Experiment 10: Cardiac fibroblasts 7 days post-MI

Mice were injected i.v. with 2.5µg anti-CD45.2-APC (Biolegend 109814) under isoflurane anesthesia 5 minutes before being euthanized by cervical dislocation. The heart cell suspensions were counted and a balanced number of cells from each sample were processed for further analysis. Dead cells were removed with magnetic Dead Cell Removal kit (Milteny Biotec, 130-090-101) and LS-Columns (Milteny Biotec, 130-042-901) according to the manufacturer’s instructions. Afterwards, the cells were washed with PBS+1%FCS and centrifuged at 400g for 5 minutes at 4°C. The supernatant was removed and the pellet was resuspended in TruStain FcX (Biolegend, 101320, 10µg/ml) in PBS+1%FCS, incubated for 10 minutes, and labeled with anti-MEFKS4-APC (Miltenyi Biotec130-120-802, 1:50), anti-CD45.2-Alexa700 (Biolegend, 109821, 1:200), anti-CD31-PE (BioLegend 160204, 1:100) antibodies, Fixable Viability Dye e780 (Thermofisher, 65-0865-14, 1:1000), a mix of CITE-seq antibodies (Biolegend, panel in Extended table 2 CITE-seq TotalSeq-A), and each sample with its respective TotalSeq-A hashtag antibodies (see below specificities for each experiment and Extended Table 1 for references). The antibody mixes were incubated for 20 minutes at 4°C. The cells were washed and Live/CD45iv^−^/CD45^−^/CD31^−^/MEFSK4^+^ cells were sorted using a BD FACS Aria III with a 100µm nozzle in PBS+1%BSA, washed and resuspended in PBS+0.04% ultrapure BSA (ThermoFisher, AM2618) at 1000 cells/µl final concentration. Cells were loaded in the 10x Genomics Chromium using Next GEM Single Cell 3’ HT Kit v3.1 with the aim to recover 16,000 cells. Experimental groups and hashtags: male MI-operated mice (n=5, hashtag 1-5) and female MI-operated mice (n=6, hashtag 6-11)

### ScRNA-seq library generation, sequencing and pre-processing

scRNA-seq libraries were generated by using 10x Genomics Chromium Next GEM Single Cell 3′ kit or 10x Genomics Chromium Next GEM Single Cell 3′ HT Kit version (10x Genomics) and processed according to the manufacturer’s instructions. Samples were prepared for CITE-Seq/Hashing according to the manual until the cDNA amplification step in which 1 μl (0.2 μM) ADT PCR additive primer to capture Antibody-derived tags (ADTs) and 1 μl (0.1 μM) HTO PCR additive primer to capture Hashtag oligos (HTOs) were added. After cDNA amplification 60 μl (0.6x) SPRI beads (Beckman Coulter) were added to separate the supernatant fraction that contains the ADT/HTO-derived cDNAs (<180bp) and the bead fraction that contains the mRNA-derived cDNAs (>300bp). The mRNA-derived cDNAs (bead fraction) were processed following the standard 10x Genomics protocol. The ADT/HTO-derived cDNAs (supernatant fraction) were purified twice with 2x SPRI. After the two purifications, half the amount of the eluted cDNAs were amplified for ADTs and half for hashtags. In the same reactions, the ADTs were indexed using TruSeq Small RNA primers and the hashtags using modified TruSeq DNA primers. The libraries were purified once more with 1.6x SPRI (160 μl). All libraries were quantified by QubitTM 3.0 Fluometer (ThermoFisher) and quality was checked using 2100 Bioanalyzer with High Sensitivity DNA kit (Agilent). Sequencing was performed with either S1 or S2 100bp flowcell with Novaseq 6000 platform (Illumina) and the reads for CITE-Seq/Hashing sample were allocated as follows: 5% for the hashtags, 10% for the ADTs and 85% for the mRNAs. 10x Genomics data, including HTO and ADT libraries, was demultiplexed using Cell Ranger software (v. 7.0.1). Mouse mm10 reference genome was used for the alignment and counting steps. The gene-barcode matrix obtained from Cell Ranger was further analyzed using Seurat (v5.3.1).

### Single-cell RNA-seq analysis

Analysis of heart CD45^+^ cells 5 days post-surgery of mice treated with isotype or anti-Ly6G: Experiments 1, 2 and 3 were preprocessed individually to remove doublets and cells not clearly assigned to a sample based on the hashtag signal, dead cells (i.e. cells with high mitochondrial transcripts content), and assign each cell to its sample of origin according to the hashtag signal. The three experiments were then integrated with Harmony ^20^. The cell clustering was performed with the “FindClusters” function within Seurat^21^ and the resolution parameter adjusted according to the differentially expressed genes across clusters. According to the cell repartition per sample, 3 samples (indicated in Extended Table 1b) were excluded due to poor immune cell number recovery. Clustering analysis of total CD45^+^ cells was performed using 30 principal components and a 0.2 resolution for the identification of the main cell subtypes (**Supplementary Figure 4a**) and at a 0.6 resolution for the identification of subpopulations of immune cells (**Figure 5a**). Cells corresponding to neutrophils were identified, and neutrophils from MI samples were separately reclustered using 20 principal components and a 0.5 resolution.

#### Analysis of bone marrow, spleen, blood and heart cells 1 and 4 days post surgery

Experiments were preprocessed individually to remove doublets and cells not clearly assigned to a sample based on the hashtag signal, dead cells (i.e. cells with high mitochondrial transcripts content), and assign each cell to its sample of origin according to the hashtag signal. Neutrophils were identified, extracted and integrated with Harmony^20^. The cell clustering was performed with the “FindClusters” function within by Seurat^21^ and the resolution parameter adjusted according to the differentially expressed genes across clusters. 2 samples (indicated in Extended Table 1b) were excluded due to poor immune cell number recovery. Clustering analysis of total neutrophils was performed using 20 principal components and a 0.3 resolution. Heart neutrophils were extracted and reclustered using 20 principal components and 0.3 resolution.

#### Analysis of cardiac fibroblasts 7 days post MI

Data was preprocessed individually to remove doublets and cells not clearly assigned to a sample based on the hashtag signal, dead cells (i.e. cells with high mitochondrial transcripts content), and assign each cell to its sample of origin according to the hashtag signal. The cell clustering was done with the function “FindClusters” function provided by Seurat^21^ and the resolution parameter adjusted according to the differential expressed genes across clusters. Fibroblasts were extracted and integrated with Harmony ^20^ with other cell subsets to perform cell-cell interaction analysis.

#### Extended analysis

Cell clusters were identified according to top10 differentially expressed genes using “FindAllMarkers” in Seurat, and wherever applicable confirmed at the cell surface marker levels using the CITE-seq signal. Additional top 7 variables features calculated as a z-score and expression of known markers from the literature presented as enhanced dot plot were generated with SeuratExtend (v.1.2.4)^22^. Clusters distribution across organ and conditions were visualized with ClusterDistrBar from SeuratExtend. Differentially expressed genes between conditions were identified by pseudo-bulk analysis: briefly, raw counts for each gene in all cells were aggregated per sample to create a pseudo-bulk matrix. The pseudo-bulk count matrices were analyzed using DESeq2 version 1.49.4^23^ and an adjusted p value cutoff of 0.05 was applied for identification of differentially expressed genes. Differentially expressed genes lists were used to build Venn diagram with ggvenn (v. 0.1.19) by ggplot2 (v. 4.0.0) or presented as Violin Plot with EnhancedVolcano (v.1.27.0). Gene expression scores were assigned using the “AddModuleScore” function within Seurat. Gene lists used for this purpose are provided in Extended Table 4. Probability of communication pathways and signaling pathway network were identified using CellChat (v.2.2.0)^24,25^. Gene ontology analysis was performed using “Find Markers” and “enrichGO” analysis. Pseudotime analysis was performed with monocle3 (v.1.4.26)^26,27^.

### Cell staining and flow cytometry analysis

Single cell suspensions were plated in a 96 round-bottom well plate and centrifuged at 400g for 5 minutes at 4°C. Pelleted cells were resuspended in TruStain FcX™ (Biolegend, 101320, 10µg/ml) and incubated for 10 minutes. Cells were then stained with primary antibodies (Extended Table 3) and Fixable Viability Dye eFluor™ 780 (Thermofisher, 65-0865-14, 1:1000) for 20 minutes at 4°C. After washing, the cells were resuspended with PBS+1%FCS and ready for FACS measurements. Acquisition was done by FACS Celesta (BD Biosciences) and analyzed with Flowjo v10.

### Histological analyses

*Heart sections preparation:* Right after mouse sacrifice as described above, KCl was injected in the exposed heart to arrest it in diastole. The heart was then perfused with cold PBS, collected, embedded in Tissue-Tek® O.C.T. Compound (Tissue-Tek®, 4583) and snap frozen in isopentane pre-cooled in liquid nitrogen and then stored at −80°C. Heart sections were cut with Cryostat (Leica, CM3050 S) in 8µm thick sections, applied on Superfrost slides™ (Epredia, 3050-002A) and stored at −80°C for further staining.

*Sirius red staining:* Slides were incubated 10min in Xylene, rehydrated in decreasing concentrations of Ethanol, then incubated 10 min in water. The slides were then stained with Picrosirius red solution (Sigma Aldrich, Direct Red 80, 365548) for 1h, 0.5% acidified water two times 1min, then dehydrated with increasing concentrations of ethanol solutions and incubated 10min in Xylene. The slides were then mounted with Eukitt^®^ Quick-hardening mounting medium (Sigma Aldrich, 3989). Picture acquisition was done with the Leica DM 4000 B LED microscope, a first acquisition was done in bright-field to assess the infarct size, a second fluorescence acquisition was done to assess the collagen deposition (evaluation of collagen autofluorescence in red and live cells in green^28^). Infarcted myocardium size measurements and collagen quantification were performed with ImageJ software (Fiji).

*Immunofluorescent staining:* fresh frozen sections were rinsed in PBS for 5 minutes and incubated with PBS+1%BSA for 30 minutes at room temperature. Then, the sections were incubated with primary antibodies (Extended Table 3) in PBS+1%BSA overnight at 4°C. The following day, they were washed in PBS for 5 minutes and then stained with the secondary antibody (Extended Table 3) in PBS+1%BSA for 1 hour at room temperature. Following the secondary antibody incubation, the sections were washed in PBS for 5 minutes and finally sealed with mounting medium containing Hoechst. Images of CD68 staining were captured with the Olympus SLIDEVIEW™ VS200 and analyzed with Qu-Path-0.5.1, images of Ly6G and PDGFRα were captured with the Zeiss imager. Z1 ApoTome.2 and analyzed with Zeiss Zen-3.11.

### RNA isolation, cDNA generation and qPCR analysis

Total RNA was extracted from either frozen tissue sections or PBS-washed cultured cells using the NucleoSpin RNA extraction Kit (Macherey Nagel, 740955) in accordance with the manufacturer’s instructions. cDNA was synthesised with the First Strand cDNA Synthesis kit (Thermofisher, K1612). Quantitative real-time polymerase chain reaction was performed on template cDNA with either: PowerUp™ SYBR™ Green Master Mix (Applied Biosystems, A25742) analyzed with an Applied Biosystems™ QuantStudio™ 6 Flex Real-Time PCR System, or with qPCR BioSyGreen Blue Mix HiRox (PCRBiosystems, PB20.16.05) and analyzed with a Step One Plus Applied Biosystems™. Quantitative measurements were determined using the ΔΔCt method, with *Hprt* as the housekeeping gene. Primer sequences are provided in Extended Table 5.

### Co-culture and scratch assays

*γδ T cell isolation and culture:* After mouse sacrifice, the spleen or thymus were collected, mechanically dissociated, filtered with a 70µm cell strainer and washed. Spleen cells were resuspended in erythrocyte lysis buffer (150 mM Ammonium chloride NH4Cl, 10 mM Potassium bicarbonate KHCO3, and 0.1 mM EDTA in distilled water) for 10min, then washed. γδ T cells were isolated with mouse TCRγ/δ T Cell Isolation Kit (Miltenyi Biotec, 130-092-125) according to the manufacturer instructions. The purity was checked by flow cytometry analysis. γδ T cells were either used directly for further experiments in a non-stimulated state or stimulated and amplified for 5 days in complete medium (RPMI Glutamax, 10%FCS, 1%P/S, 10µM Betamercaptoethanol, 10mM HEPES, 1mM Na Pyruvate, 100µM non-essential amino acids) together with anti-CD3 (Thermofisher, 16-0032-85, 2µg/mL), anti-CD28 (Thermofisher, 16-0281-85, 2µg/ml) and Il-2 (PeproTech®, 212-12, 10ng/mL).

*Neutrophil isolation and culture:* After mouse sacrifice, bone marrow cells were collected from the hindlimb bones, filtered with 70µm cell strainer and washed once with PBS+1%FCS. Neutrophils were isolated with mouse neutrophil isolation kit (Miltenyi Biotec, 130-097-658) according to the manufacturer instructions. After cell count, neutrophils were differentiated into SiglecF^+^ neutrophils by incubation of 2.10^6^ neutrophils per ml of medium (RPMI Glutamax, 10%FCS, 1%P/S, 10µM Betamercaptoethanol) supplemented with recombinant Mouse GM-CSF (PeproTech®, 315-03, 10ng/ml) and recombinant Mouse TGF-β1 (Biolegend, 763102, 10ng/ml) for 48h. Before further use, SiglecF^+^ neutrophils were washed twice in medium.

*Co-culture with fibroblasts and scratch assay:* NIH 3T3 Fibroblast (ATCC, CRL-1658 ^™^) cells were plated in cell culture plate and left overnight to attach to reach around 60% confluence (250 000 cells in P12 plate). For quantitative PCR experiments: The next day, γδ T cells were added on top of the adherent fibroblasts at a ratio 1 fibroblast for 1 γδ T cell and co-cultured in starving medium (DMEM Glutamax, 1%FCS, 1%P/S) for 24, 48 or 72h. At the end of the experiment, non-adherent γδ T cells were washed away with PBS and remaining adherent fibroblasts were processed for RNA extraction. For scratch assay experiments: the fibroblast layer was scratched with the help of a 200µl tip. The medium and detached cells were removed and fresh starving medium (DMEM Glutamax, 1%FCS, 1%P/S) was added. Depending on the experimental condition, 400 000 γδ T cells or 1 200 000 neutrophils were added, and additionally treated with: 0.1% DMSO in control wells when applied, 10µM Cyclophilin A inhibitor (Sigma-Aldrich, 239836, diluted in DMSO), 10µM Galunisertib (Sigma-Aldrich, SML2851, diluted in DMSO). Images were acquired every hour in bright field with a Nikon Eclipse Ti and analyzed with ImageJ-1.54g. Quantification of wound closure was performed with ImageJ software (Fiji).

### Statistical analysis

Statistical analysis was performed with Graphpad Prism Software 10. Unpaired t-test was used for 2 groups comparison. Ordinary one-way ANOVA (with a Fisher’s LSD test to compare specific experimental groups) was used for more than 2 groups comparison. The data are presented as mean ± SEM and p values<0.05 were considered statistically significant.

### Data availability

Single-cell RNA-sequencing data generated for this report will be made available upon publication.

## Results

### Myocardial infarction transcriptionally primes neutrophils in the periphery

To characterize the landscape of neutrophil heterogeneity and recruitment kinetics following myocardial infarction (MI), we performed single-cell RNA sequencing on neutrophils harvested from the bone marrow, spleen, blood, and heart. Samples were collected at days 1 and 4 post-permanent MI—as well as from control and sham-operated mice—to capture the full trajectory from the initial peak of cardiac infiltration (day 1) to the emergence of the SiglecF^+^ population (day 4) (**Figure 1a**)^13,16,29^. In peripheral organs, we observed a continuum of neutrophil states, from progenitors and pre-neutrophils towards mature neutrophils (**Figure 1a-d, Supplementary Figure 1a,b**). We recovered progenitors and pre-neutrophils (prog/pre-Neu) expressing high levels of proliferating genes, immature neutrophils (*Ltf*, *Camp*, *Ngp*, *Lcn2*), young neutrophils (*Mmp8, Prok2* and *Retnlg*), two subsets of mature neutrophils (‘Mature 1’ and ‘Mature 2’) expressing high levels of *Ccl6* and *Cxcr2*, one cluster with a type I interferon response signature (IFN; *Isg15*, *Ifit3*), and a small cluster of *Cd244a*^+^ neutrophils. All these subsets were found in the bone marrow and spleen, while only few young neutrophils were found in the blood, together with mature and IFN subsets (**Figure 1e, Supplementary Figure 1c**). Finally, a heart-specific cluster displayed high expression of activation markers (e.g. *Cxcl2*, *Ccl3*, *Ccl4*, *Il1a*, *Il23a*) but also markers specific for previously described SiglecF^+^ neutrophils^16^ such as *Icam1*, *Siglecf*, and enrichment in anti-apoptotic genes including *Bcl2a1d* (**Figure 1b,c, Supplementary Figure 1b**).

**Figure 1:**
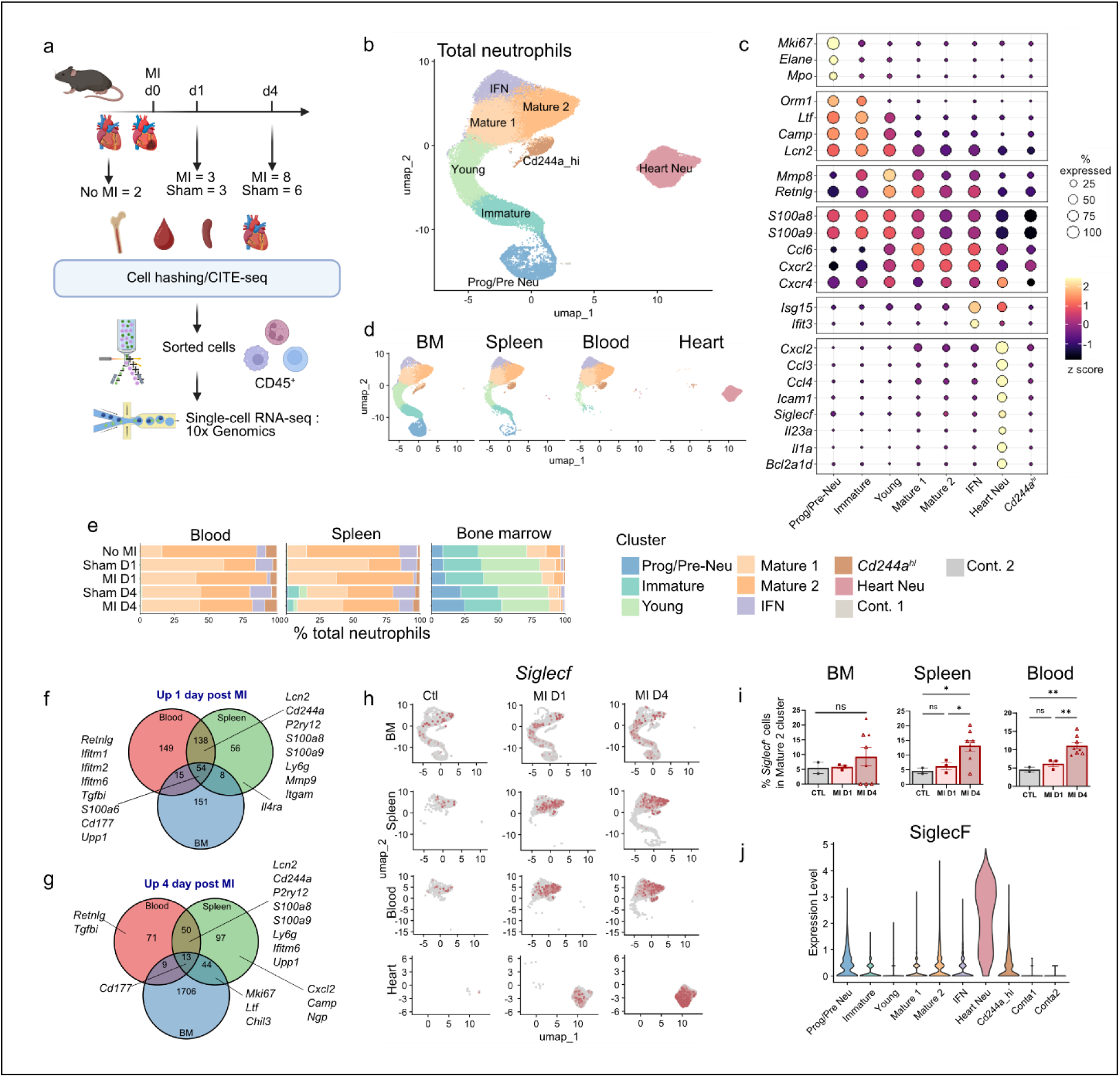
Neutrophils are primed in hematopoietic organs after myocardial infarction. **a)** Experimental design of the single cell RNA-seq/CITE-seq experiments. **b)** Integrated UMAP plot visualization of neutrophils extracted from each organ and timepoints. **c)** Relative expression of the main marker genes used to identify neutrophil clusters in (b). **d)** UMAP plot visualization of neutrophils according to organ of origin. **e)** Neutrophil clusters proportions in blood, spleen and bone marrow across conditions. **f,g)** Venn diagram showing upregulated genes in 1- and 4-days post-MI total neutrophils in the bone marrow, spleen and blood identified by pseudobulk analysis (Supplementary Figure 1d). **h)** *Siglecf* expression projected on the UMAP plot of total neutrophils (panel b) split by organ and time point. **i)** Percentage of cells with detectable *Siglecf* transcripts in the Mature-2 cluster in bone marrow, spleen, and blood in the indicated experimental conditions. One point represents one mouse, open triangles represent female mice. **j)** CITE-seq signal for SiglecF protein surface expression in the indicated neutrophil populations at day 4 after MI. MI: myocardial infarction, BM: bone marrow. Illustration (panel a) created with Biorender. Statistical tests (panel i): Ordinary one-way ANOVA (Fisher’s LSD test) (*<0.05; **<0.01; ***<0.001; ****<0.0001).

To evaluate how MI globally affected peripheral neutrophils, we performed differential pseudo-bulk gene expression analysis in total neutrophils (**Figure 1f,g, Supplementary Figure 1d**), and differential neutrophil subset composition (**Figure 1e, Supplementary Figure 1e**) analysis across experimental groups. Neutrophil gene expression profiles were affected in all compartments at 1 and 4 days after MI, with specific patterns of shared transcriptional priming across organs and time points (**Figure 1f,g, Supplementary Figure 1d**). Of note, activation markers including *S100a8/9* were upregulated in blood and spleen at both timepoints, as well as *Retnlg* at day 1 in all compartments. The *Cd177* marker, associated with neutrophil activation and endothelial transmigration^30–32^, was also upregulated in all compartments and at all timepoints after MI. The percentage of prog/pre-Neu was increased in the bone marrow 4 days after MI. The spleen also displayed appearance of prog/pre-Neu, immature and young neutrophils at day 4 (**Figure 1e, Supplementary Figure 1e)**, suggesting an increased granulopoiesis in both hematopoietic compartments. Consistently, we observed an upregulation of genes associated with proliferation and the pre-neutrophil state, such as *Mki67* and *Ltf*, in the bone marrow and spleen 4 days post-MI. Furthermore, markers of pre- and immature neutrophils, including *Camp* and *Ngp*, were specifically enriched in the splenic neutrophil population at this same time point. *Upp1* expression, a gene known to be upregulated during emergency granulopoiesis^33,34^, was increased in all compartments at day 1, and was still upregulated at day 4 in the spleen and blood, suggesting a prolonged granulopoiesis in the spleen following MI (**Figure 1f,g**). Altogether, these data indicate that MI triggers granulopoiesis and transcriptional priming of neutrophils in the periphery.

Using cellular indexing of transcriptomes and epitopes by sequencing (CITE-seq) ^35^ in infarcted mice, we have previously shown that neutrophils acquired surface protein expression of SiglecF only in the ischemic heart tissue, and not in the blood^16^. Nevertheless, previous evidence suggests a potential for peripheral maturation, where circulating neutrophils are primed to express *Siglecf* transcripts prior to extravasation, as observed in lung cancer settings^19^. To specifically assess whether the acquisition of the SiglecF^+^ state could start in the periphery after MI, we first analyzed *Siglecf* expression in all recovered neutrophil populations (**Figure 1h**). As expected, *Siglecf* expression was high in heart neutrophils at day 4 post-MI. Nevertheless, we detected *Siglecf^+^*neutrophils within the ‘mature 2’ cluster (**Figure 1h**), with an increased percentage of cells with detectable *Siglecf* transcripts in spleen and blood ‘mature 2’ neutrophils at 4 days post-MI (**Figure 1i**). Consistent with previous observations, SiglecF surface protein measured by CITE-seq was only substantially expressed in heart neutrophils at day 4 (**Figure 1j**). Altogether, these results suggest that MI can transcriptionally prime neutrophils in the periphery towards the SiglecF^+^ signature, but that they only fully acquire it once in the ischemic heart tissue.

### GM-CSF and TGFβ drive the ischemic heart-acquired SiglecF^+^ neutrophil signature

We next explored the cardiac neutrophil population. Heart neutrophils were extracted and reclustered. We identified five distinct clusters: ‘mature’ neutrophils, ‘mature Tlr4^hi^’ neutrophils presenting high expression level of *Tlr4* and *Cxcl3*, an IFN cluster expressing interferon inducible genes, and two SiglecF^+^ clusters: SiglecF^+^Tnf^hi^ expressed high level of *Tnf* but also markers previously associated to the SiglecF^+^ signature^16^, and SiglecF^+^ neutrophils enriched in *Siglecf* and *Cxcr4* (**Figure 2a, Supplementary Figure 2a,b**). Each subset displayed specific putative functions and activities, and a time-dependent appearance. The ‘mature Tlr4^hi^’ subset was enriched for gene ontology terms suggesting a strong response to hypoxia environment and displayed a high NETosis score, in line with its prevalence at the early day 1 timepoint. The ‘mature’ cluster, present at both day 1 and day 4 post-MI, was involved in leukocyte chemotaxis and cell migration processes. Both SiglecF^+^ subsets were prevalent at day 4: the SiglecF^+^ cluster was enriched for pathways associated with RNA splicing and exit from the nucleus, while the SiglecF^+^Tnf^hi^ subset was enriched for ribosomal activities and ROS production (**Figure 2 b,c, Supplementary Figure 2c,d**). Altogether, these data are consistent with previous work showing time-dependent apparition of ROS-producing SiglecF^+^ neutrophils in the ischemic heart^13,16^.

**Figure 2:**
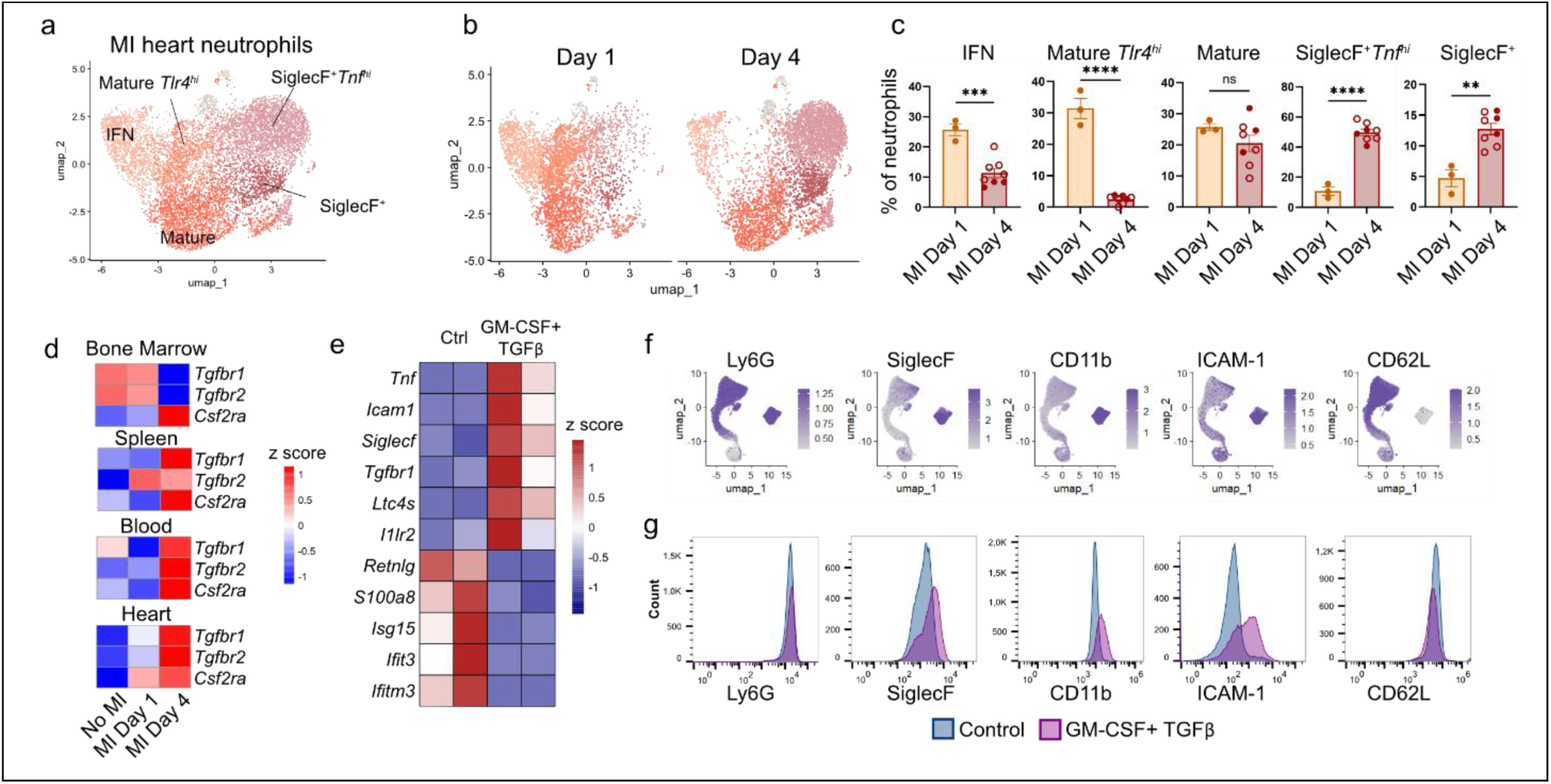
Terminal differentiation of SiglecF^+^ neutrophils occurs in the infarcted heart. **a)** UMAP plot visualization of heart neutrophils extracted from Figure 1b and reclustered. **b)** UMAP plot from panel a) split by timepoint after MI. **c)** Percentage of heart neutrophil clusters in total heart neutrophils according to time post MI. One circle represents one mouse, open circles represent female mice. **d)** Relative expression of *Csf2ra*, *Tgfbr1* and *Tgfbr2* transcripts in total neutrophils in the indicated conditions. **e)** relative gene expression in neutrophils left untreated or GM-CSF+TGFβ stimulated neutrophils assessed by quantitative PCR (n= neutrophils from 2 mice). **f)** CITE-seq signal of the indicated surface protein on bone marrow, spleen, blood and heart neutrophils at day 4 post-MI projected onto the UMAP plot from Figure 1b. **g)** Surface protein expression of the indicated markers on pre-gated live Ly6G+ neutrophils assessed by flow cytometry, in unstimulated bone marrow-isolated neutrophil, and GM-CSF+TGFβ stimulated neutrophils. MI: myocardial infarction, BM: bone marrow, CTL: control. Statistical tests: (panel c) unpaired t-test (*<0.05; **<0.01; ***<0.001; ****<0.0001).

Besides MI, SiglecF^+^ neutrophils were described in chronic kidney disease associated with kidney fibrosis, in which they were found to be derived from GM-CSF- and TGFβ1-stimulated conventional neutrophils^17^. Therefore, we examined the expression of GM-CSF and TGFβ receptors in post-MI neutrophils. *Csf2ra, Tgfbr1*, and *Tgfbr2* were highly expressed after MI in spleen and blood neutrophils and further increased over time post-MI, although this upregulation was not observed in the bone marrow. In particular, cardiac neutrophils subsets upregulated *Tgfbr1*, *Tgfbr2*, and *Csf2ra* receptors over time post-MI. (**Figure 2d**), suggesting that TGFβ, together with GM-CSF signaling activation in neutrophils, could fuel the acquisition of the SiglecF^+^ state. Exposure of bone marrow-isolated neutrophils to a combination of GM-CSF and TGFβ induced the SiglecF^+^ signature both at the transcriptomic level with upregulation of *Siglecf*, *Tnf*, *Icam1*, *Tgfbr1*, *Ltc4s* and *Il1r2* and downregulation of *Retnlg*, *S100a8*, *Isg15*, *Ifit3* and *Ifitm3* (**Figure 2e**), and at the protein level, recapitulating the cell surface protein signature of cardiac SiglecF^+^ neutrophils with increased SiglecF, ICAM-1, and CD11b, and decreased CD62L (**Figure 2 f,g**). Altogether, these results suggest that cardiac SiglecF^+^ neutrophil terminal differentiation is driven by GM-CSF and TGFβ from peripherally primed neutrophils post-MI.

### Ly6G targeting increases fibrosis and transcriptionally reprograms cardiac neutrophils after MI

A previous study employing Ly6G targeting showed a detrimental effect on post-MI remodeling caused by defects in macrophage polarization^5^. However, we have previously demonstrated that the administration of the rat anti-mouse Ly6G antibody (clone 1A8) fails to effectively deplete the cardiac neutrophil pool at day 3 post-MI. Instead, this intervention triggers a phenotypic reprogramming of the remaining cells toward a SiglecF^+^ state^16^. These findings imply that the adverse effects of anti-Ly6G treatment are driven by a shift toward a SiglecF^+^ neutrophil identity rather than by successful neutrophil depletion. In an attempt to strengthen the efficiency of neutrophil depletion, we used a combination of rat IgG2aκ anti-Ly6G and a secondary mouse anti-rat IgG2aκ^36^ as a Ly6G targeting strategy. The mice were randomized into two groups, a rat IgG2aκ isotype control + mouse anti-rat IgG2aκ treated group (Isotype group) and a rat IgG2aκ anti-Ly6G clone 1A8 + mouse anti-rat IgG2aκ treated group (anti-Ly6G group). The mice were treated daily starting 2 days before MI induction and until 7 days post-MI (**Figure 3a**). Despite using this novel strategy to target neutrophils, we observed similar results as in our previous study^16^: at day 3 post-MI, cardiac neutrophil levels were only decreased by ∼40% in the anti-Ly6G group, which furthermore presented a shift toward SiglecF^+^ neutrophils (**Figure 3b**). Ly6G targeting failed to influence survival or infarct size but substantially increased border-zone fibrosis (**Supplementary Figure 3a-b** and **Figure 3c**), echoing the pro-fibrotic impact observed in earlier studies ^5^. Consequently, we leveraged this treatment as a targeted approach to uncover how shifts in neutrophil programming drive the functional and structural deterioration of the infarcted myocardium.

**Figure 3:**
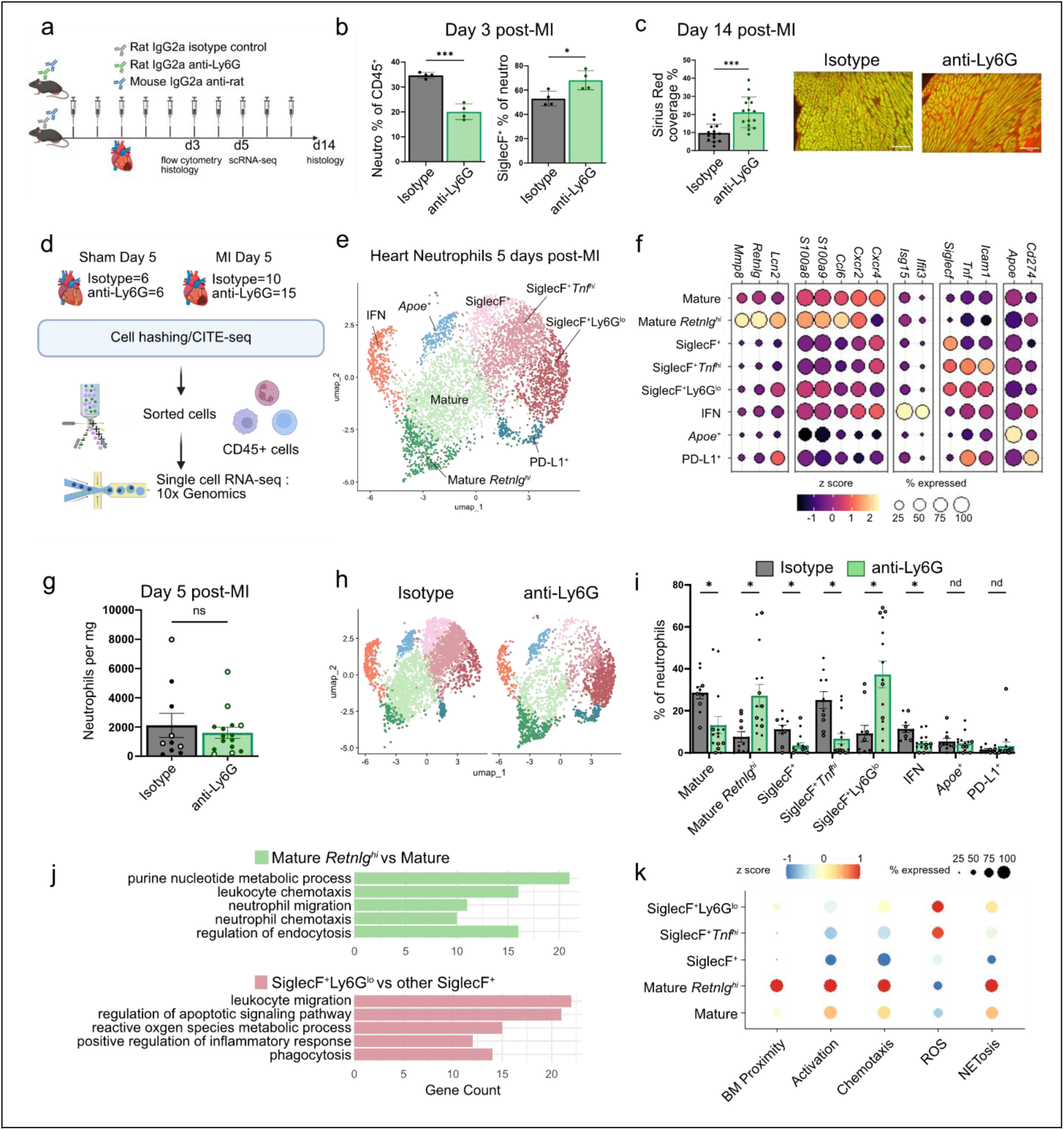
Anti-Ly6G reprograms heart neutrophils after myocardial infarction. **a)** Experimental design of Ly6G targeting experiments. **b)** Flow cytometry analysis of heart neutrophils 3 days post MI: percentage of Neutrophils among total CD45^+^ cells (left panel) and percentage of SiglecF^+^ neutrophils among total neutrophils (right panel). **c)** Interstitial fibrosis in the infarct border zone 14 days post-MI, expressed as percentage of total area, determined by picrosirius red staining, scale bar: 100µm. **d)** Experimental design of the single cell RNA-seq/CITE-seq experiments with Ly6G targeting. Group isotype MI n= 4 females and 6 males, group anti-Ly6G MI n= 5 females and 10 males, group isotype sham n= 3 males and 3 females, group anti-Ly6G sham n= 3 males and 3 females. **e)** UMAP plot of neutrophils identified in (Supplementary Figure 4a), extracted from the MI groups and reclustered. N=3 independent scRNA-seq/CITE-seq experiments were integrated. **f)** Relative expression of the main marker genes used to identify neutrophil subsets. **g)** Neutrophil counts per mg of heart tissue at day 5 post-MI. **h)** UMAP plot of neutrophils from panel e) split by treatment. **i)** Neutrophil cluster proportions expressed as percentage of total neutrophils. **j)** Gene count for 5 of the top16 up regulated biological processes in the Mature *Retnlg^hi^* cluster vs. the Mature cluster (green bars) and in the SiglecF^+^Ly6G^lo^ cluster vs. the all SiglecF+ neutrophil clusters (pink bars); all with adjusted p-value <0.05 **k)** Expression of the indicated scores by neutrophil clusters from panel e). Bar plots: one circle represents one mouse, open circles represent female mice. MI: myocardial infarction, BM: bone marrow, Iso: isotype, GO: gene ontology. Statistical tests: (panel b, c, g, i) unpaired t-test (*<0.05; **<0.01; ***<0.001; ****<0.0001). Illustration (panel a,d) created with Biorender.

To further characterize neutrophils in the infarcted heart after targeting Ly6G, and possible associated effects on the cardiac immune landscape, such as macrophage phenotypic shifts^5^, we performed single cell RNA-seq/CITE-seq analysis of cardiac CD45^+^ cells 5 days after MI or sham surgery, in isotype or anti Ly6G-treated mice (**Figure 3d**). Importantly, cardiac neutrophil counts remained comparable between groups at this time point, signifying a total loss of the initial depletion effect and confirming that the observed phenotypes were not due to numerical differences in the infiltrate (**Figure 3g**). From total CD45^+^ cells (**Supplementary Figure 4a,b**), we extracted neutrophils from the MI groups and performed a high-resolution clustering to distinguish neutrophil subpopulations (**Figure 3e**). Transcriptomic and protein surface markers analysis revealed 8 neutrophil subsets: mature neutrophils, mature-*Retnlg^hi^* neutrophils, three SiglecF^+^ neutrophil clusters, including a SiglecF^+^Ly6G^lo^ cluster, an IFN response cluster, and two minor clusters defined as *Apoe*^+^ and PD-L1^+^ (**Figure 3f, Supplementary Figure 4c,d)**.

Interestingly, the mature-*Retnlg*^hi^ and the SiglecF^+^Ly6G^lo^ clusters were mostly specific to the anti-Ly6G group, while mature, SiglecF^+^ and SiglecF^+^*Tnf*^hi^ clusters were more represented in the isotype group (**Figure 3h,i**). Mature-*Retnlg*^hi^ neutrophils highly expressed genes usually related to ‘young’ and activated neutrophils, including *Retnlg*, *Ccl6*, and *Mmp8* (**Figure 3f**). The two neutrophil subsets observed specifically after Ly6G-targeting at day 5 post-MI, mature-*Retnlg*^hi^ and SiglecF^+^Ly6G^lo^, revealed enrichment in pathways and gene expression involved in leukocyte chemotaxis and migration, and in NETosis, compared to control-associated counterparts (**Figure 3 j,k**). The mature-*Retnlg*^hi^ cluster further displayed enhanced neutrophil migration and chemotaxis enrichment and a high bone marrow proximity score, while the SiglecF^+^Ly6G^lo^ cluster increased regulation of apoptotic signaling pathways, inflammatory response, phagocytosis, and ROS metabolic processes (**Figure 3j,k**). Total CD45^+^ cells in the bone marrow were increased in the anti-Ly6G group (**Supplementary Figure 4e**) suggesting enhanced hematopoiesis. Altogether, these results indicate that Ly6G targeting does not efficiently deplete cardiac neutrophils after MI but induces a shift towards SiglecF^+^ neutrophils at day 3 after MI, and complex modifications of the neutrophil landscape at day 5 with the emergence of discrete populations of SiglecF^+^Ly6G^lo^ and mature-*Retnlg*^hi^ neutrophils, and ultimately results in increased fibrosis.

### SiglecF^+^ neutrophils interact with fibroblasts to enhance fibrosis through TGFβ and CypA signaling

Consequently, we sought to investigate the pathways by which the shift in neutrophil identity promotes fibrotic remodeling, exploring how these programmed changes drive the structural deterioration of the infarcted heart. Recent evidence suggests that neutrophils express collagen and can be directly responsible for collagen deposition in injured tissues^17,37^.

To test this possibility in the MI context, we applied extracellular matrix (ECM) related, collagen, and secreted factors scores extracted from the molecular signatures database (MSigDB) of the Matrisome project^38,39^ to an in-house scRNA-seq atlas of post-MI non-cardiomyocyte myocardial cells. Post-MI cardiac neutrophils did not present any expression of scores related to ECM glycoproteins, ECM regulators, proteoglycans or collagens, but presented a slight expression for ECM affiliated proteins, in contrast to fibroblasts and myofibroblasts, which highly expressed all these matrisome components (**Figure 4a**). However, neutrophils presented the highest score for matrisome-related secreted factors (**Figure 4a**), suggesting that shifts in neutrophil populations could affect fibrosis through paracrine interactions with other myocardial cells. Three days after MI, neutrophils and fibroblasts localized in close proximity in the border zone of the infarcted region (**Figure 4b**), suggesting a possible direct interaction between neutrophils and fibroblasts. Furthermore, we observed increased labeling for PDGFRα, a general fibroblast marker, in the border zone of the infarcted area of anti-Ly6G-treated mice at this time point (**Figure 4c**). Fibroblasts co-cultured with SiglecF^+^ neutrophils displayed higher capacities of wound closure in a scratch assay compared to control fibroblasts (**Figure 4d**). Cellular crosstalk analysis with CellChat ^24^ between heart neutrophils (**from Figure 2a**) and cardiac fibroblasts identified specific interactions between SiglecF^+^ neutrophils and fibroblasts, that could be responsible for fibroblast proliferation and activation, and thus fibrogenic activity (**Figure 4e**). In addition to tumor necrosis factor (TNF) signaling pathways, already described as a potential mediator of fibroblast activation by SiglecF^+^ neutrophil^40^, CypA (or Cyclophilin A, encoded by *Ppia*) and TGFβ-related (*Tgfb1*-*Tgfbr1/2* and *Tgfb1-Acvr1*) signaling pathways appeared to be the most likely involved in SiglecF^+^ neutrophil crosstalk with fibroblasts (**Figure 4e,f**). Fibroblasts incubated with SiglecF^+^ neutrophils together with Galunisertib, a TGFβ receptor kinase inhibitor (**Figure 4g**), or with a Cyclophilin A inhibitor (**Figure 4h**), presented decreased wound closure capacities compared to SiglecF^+^ neutrophil incubation alone, suggesting that SiglecF^+^ neutrophils enhance fibroblast migration and scar formation through TGFβ and Cyclophilin A signaling.

**Figure 4:**
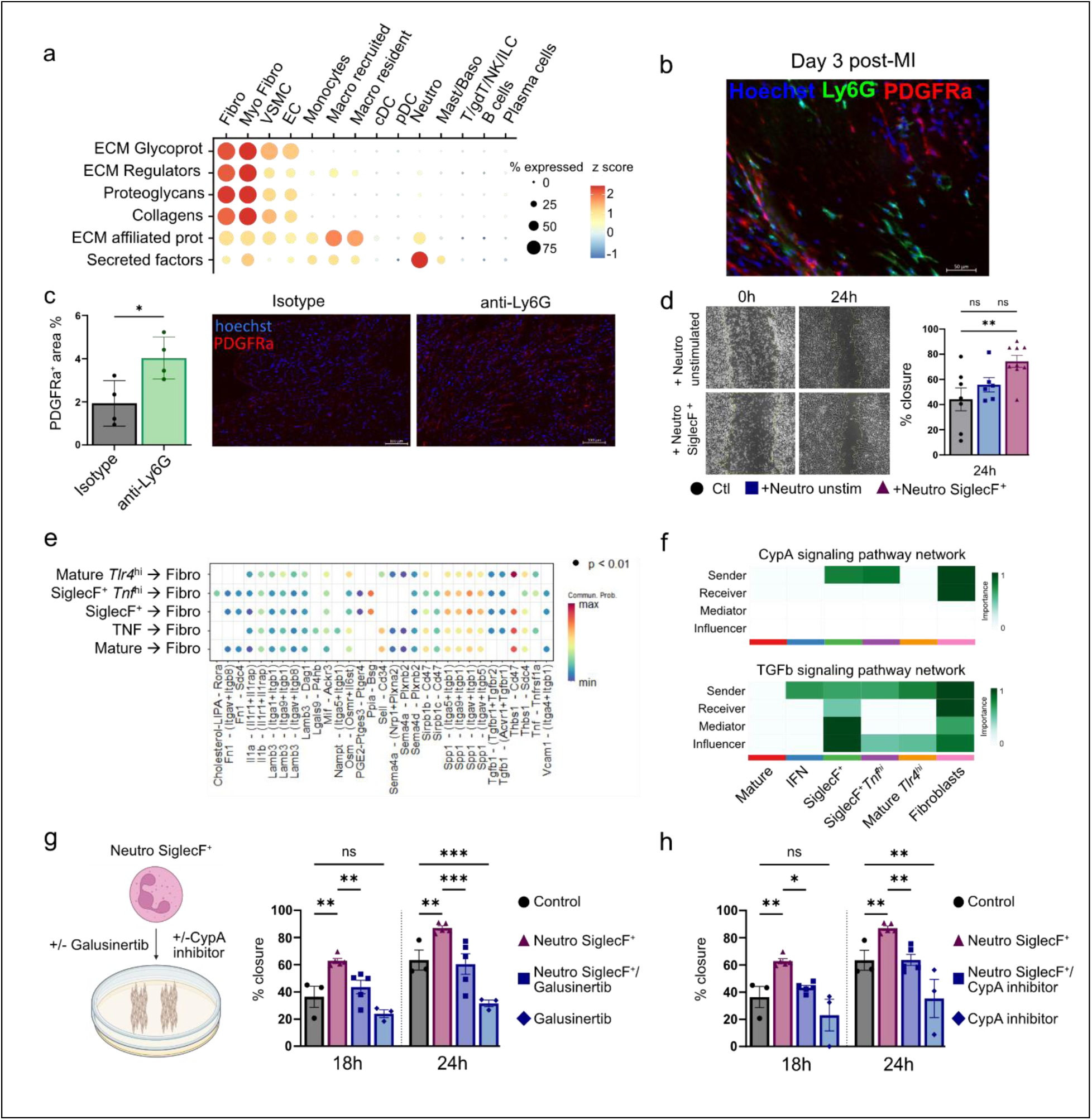
SiglecF^+^ neutrophils can interact with fibroblasts to enhance fibrosis. **a)** Expression of the indicated scores in control and day 5, 7 and 14 post-MI heart cells from an in-house scRNA-seq atlas. **b)** Immunofluorescent staining of day 3 post-MI heart section showing the infarct and border zone area. Hoechst is stained in blue, Ly6G (neutrophils) in green and PDGFRα (fibroblasts) in red, scale bar: 50µm. **c)** Immunofluorescence staining of PDGFRα in the peri infarct region 3 days post-MI, expressed as PDGFRα^+^ area percentage of total area (left panel). PDGFRα is stained in red and Hoechst in blue (right panel), scale bar: 100µm. **d)** Scratch assay on 3T3 fibroblasts incubated 24h with untreated or SiglecF^+^ neutrophils; representative images at t=0 and t=24h (left panel), quantification of percentage of wound closure after 24h (right panel); each point represents one well, pooled from three independent experiments. **e,f)** CellChat analysis of heart neutrophil cluster (from Figure 2a) communication to cardiac fibroblasts; e) Probability of communication pathways (ligand-receptor pairs); f) associated CypA and TGFβ signaling pathway network. **g,h)** Percentage of wound closure after 18h and 24h from scratch assay of 3T3 fibroblasts incubated 24h with SiglecF^+^ neutrophils and Galunisertib g) or CypA inhibitor h). Each point represents one well, pooled from two independent experiments. MI: myocardial infarction, Fibro: Fibroblasts, Myo Fibro: Myofibroblatsts, VSMC: Vascular smooth muscle cell, EC: Endothelial cell, Macro: Macrophage, cDC: conventional dendritic cell, pDC: plasmacytoid dendritic cell, Neutro: Neutrophil, Mast: Mastocyte, Baso: Basophil, T: T cell, gdT: gamma delta T cell, NK: Natural Killer, ILC: innate lymphoid cell, ECM: extracellular matrix, Glycoprot: glycoproteins, prot: proteins. Statistical tests: (panel c) unpaired t-test, (panel d,g,h) Ordinary one-way ANOVA (Fisher’s LSD test) (*<0.05; **<0.01; ***<0.001).

### Targeting Ly6G modifies the immune cell landscape of the infarcted heart

The two neutrophil subsets observed specifically after Ly6G-targeting at day 5 post-MI, mature-*Retnlg*^hi^ and SiglecF^+^Ly6G^lo^ neutrophils, revealed enrichment in pathways involved in immune cells chemotaxis, migration, and activation compared to their associated counterparts in control MI mice (**Figure 3j,k**). We therefore hypothesized that anti-Ly6G-associated neutrophils could modulate immune cell infiltration in the heart. We took advantage of our day 5 single-cell RNA-seq dataset (**Figure 3d**) and reclustered post-MI cardiac CD45^+^ cells (**Supplementary Figure 4a)** to gain granularity in leukocyte subpopulations (**Figure 5a**).

**Figure 5:**
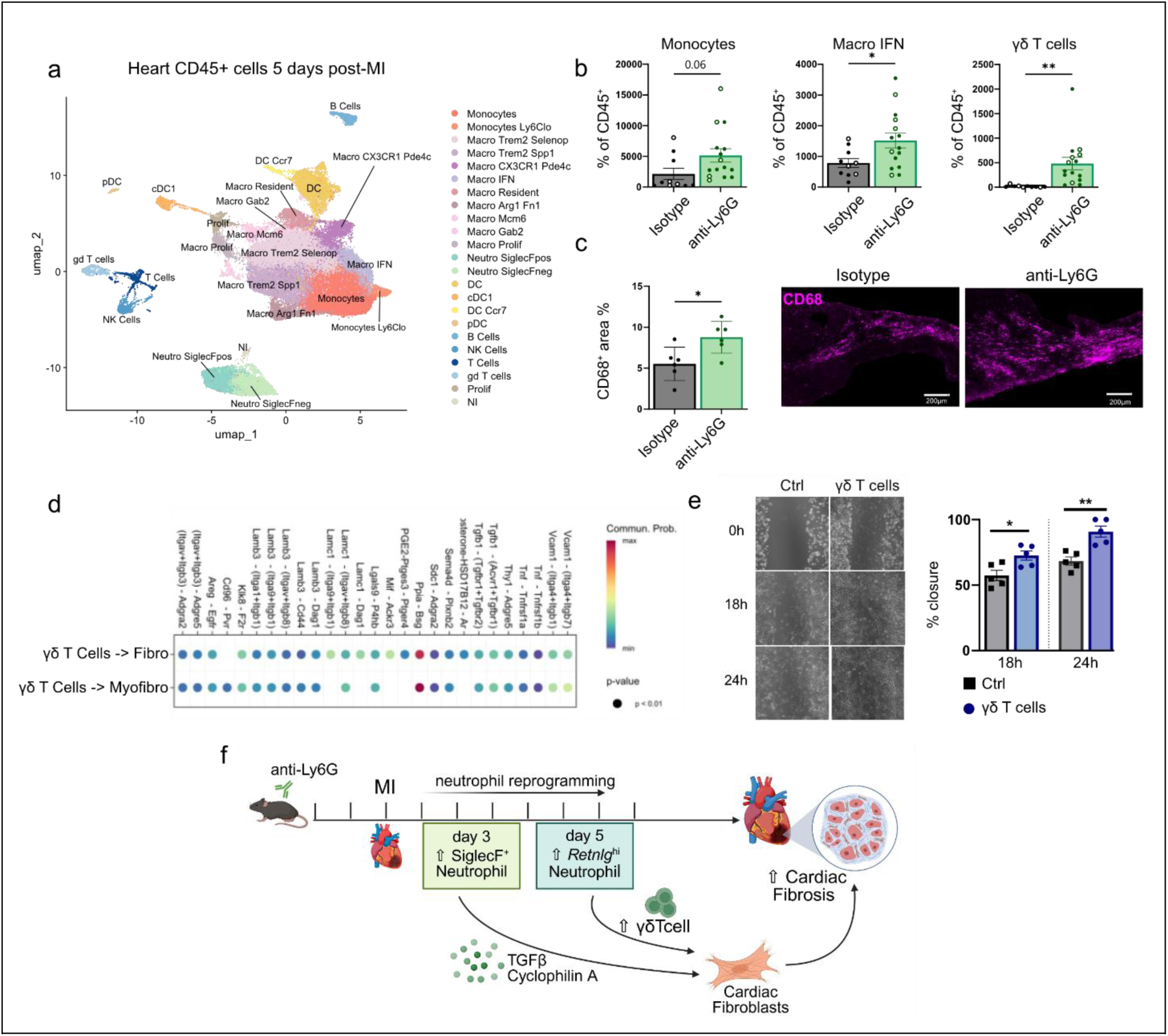
Targeting Ly6G modulates the immune cell landscape in the heart after MI with increased pro-fibrotic γδ T cells. **a)** UMAP visualization of total heart CD45^+^ cells 5 days post MI, in mice with or without Ly6G targeting (see Figure 3d). **b)** Cell count per mg of heart tissue for the indicated cell type clusters identified in (a). **c)** Macrophage content in the infarct area 14 days post-MI in isotype or anti-Ly6G treated groups. Left panel: immunofluorescence staining of CD68 (pink) on heart sections, scale bar = 200µm. Right panel: quantification of macrophage area expressed as percentage of the infarcted area. **d)** CellChat analysis of γδ T cells communication with cardiac fibroblasts, probability of communication pathways (ligand-receptor pairs) **e)** Scratch assay of 3T3 fibroblasts incubated 24h with activated γδ T cells, representative images at t=0, t=18h and t=24h (left panel) and quantification of percentage of wound closure after 18h and 24h (right panel). Each point represents one well, pooled from three independent experiments. **f)** Proposed model and summary of the main findings. Bar plots: one circle represents one mouse, open circles represent female mice. MI: myocardial infarction, CTL: control, Macro: Macrophages, Prolif: Proliferating, Neutro: Neutrophils, DC: Dendritic Cells, cDC1: conventional Dendritic Cell 1, DC: Dendritic Cells, pDC: plasmacytoid Dendritic Cells, NI: non-identified. Statistical tests: (panel b,c,e) unpaired t-test (*<0.05; **<0.01). Illustration (panel f) created with Biorender.

After MI, the anti-Ly6G group presented higher number of conventional monocytes (Monocytes) and Ly6C^lo^ monocytes, IFN macrophages and *Mcm6^+^* macrophages, and a decrease in NK cell numbers, compared to the isotype group (**Figure 5b, Supplementary Figure 5a**). Macrophage content was higher at day 14 post-MI in the anti-Ly6G group (**Figure 5c**), corroborating increased monocyte infiltration. Moreover, we observed that γδ T cell counts were drastically increased in the anti-Ly6G group (**Figure 5b**). To understand how neutrophil modulation by anti-Ly6G affects immune cell recruitment and phenotype in the heart, we ran CellChat^24^ analysis between cardiac immune cell subsets. To identify the drivers of post-MI inflammation, we screened intercellular communication pathways and identified the CCL chemokine axis as a primary candidate for orchestrating neutrophil-dependent recruitment of secondary immune populations to the heart. Refining the analysis, we showed that the most probable ligand-receptor couple involved in the recruitment of Monocytes, Macro IFN and γδ T cells was the *Ccl6*-*Ccr2* couple, with a high and specific expression of *Ccl6* by mature-*Retnlg^hi^* among all neutrophils, and high expression of *Ccr2* by the two subsets of monocytes, Macro IFN and γδ T cells (**Supplementary Figure 5b,c**), in line with the observed increased infiltration of these cells in the anti-Ly6G group. Moreover, *Ccl6* expression was increased in the cardiac tissue 3 days post-MI in the anti-Ly6G group (**Supplementary Figure 5d**). These data suggest that Ly6G targeting induces a specific population of mature*-Retnlg^hi^*neutrophils infiltrating the heart that can increase monocyte, IFN macrophage, and γδ T cell recruitment, potentially via CCL6-CCR2 signaling.

Given the drastic increase in γδ T cells in the heart following Ly6G targeting, and as they were previously described as mediators of adverse cardiac remodeling after MI^41^, we investigated their potential involvement in fibrosis. CellChat analysis revealed crosstalk between γδ T cells and fibroblasts with probable pathways implicated in the communication of γδ T cells as senders and fibroblasts as receivers (**Figure 5d**). Of note, *in silico* analysis suggested that γδ T cells are part of the sender cells in pathways involved in fibroblast proliferation and fibrosis induction including the Klk8 (encoding Kallikrein-8)/F2r (encoding Protease Activated Receptor-1, PAR1) pathway^42–45^. To functionally confirm this *in vitro*, we isolated γδ T cells from mice (**Supplementary Figure 6a**) and kept them either untreated or activated with IL-2, anti-CD3, and anti-CD28 antibodies. We then co-cultured γδ T cells with fibroblasts for 24, 48, or 72h and analyzed fibroblast gene expression. Activated γδ T cells induced expression of the proliferation-associated genes *Mki67* and *Pcna* in fibroblasts, as well as expression of the myofibroblast marker *Acta2* after 72h of co-culture, and inflammatory factors such as *Il6* and *Ccl2*, while non-activated γδ T cells did not have any effect (**Supplementary Figure 6b**). Fibroblasts co-cultured with activated γδ T cells showed increased migration in a scratch assay (**Figure 5e**), further indicating that the accumulation of γδ T cells in the heart after Ly6G targeting could be responsible for increased fibrosis. Altogether, our data suggest that targeting Ly6G shifts neutrophils towards a SiglecF^+^ state at day 3 post-MI, and induces the emergence of mature-*Retnlg^hi^* neutrophils at day 5. SiglecF^+^ neutrophils may increase fibrosis by acting on fibroblasts through TGFβ and Cyclophilin A, while at later time points, mature-*Retnlg^hi^*neutrophils may participate in the recruitment of γδ T cells which themselves contribute to enhanced cardiac fibrosis (**Figure 5f**).

## Discussion

MI precipitates a massive and rapid influx of neutrophils into the ischemic myocardium, marking the initiation of a complex inflammatory cascade. We show that these infiltrating neutrophils are not a uniform population but rather exhibit distinct temporal programming, characterized by the emergence of heterogeneous subsets such as the SiglecF^+^ cluster. Despite their abundance, the mechanisms governing this neutrophil diversification and its subsequent impact on the trajectory of cardiac repair have, until now, remained largely enigmatic. We show here that MI primes neutrophils in the bone marrow and spleen, presenting up-regulation of genes associated with neutrophil maturation, activation and migration, as well as emergency granulopoiesis-associated genes. Granulopoiesis was enhanced at day 4 with appearance of genes involved in granulocyte proliferation in both spleen and bone marrow, suggesting a replenishment of the granulocyte progenitor compartment. Interestingly, we observed an up-regulation of *Upp1* in neutrophils after MI in mice^33,34^. A recent study from Aa et al. showed increased UPP1 enzyme in the plasma of MI patients compared to control patients, further suggesting that UPP1 could originate from circulating neutrophils^46^. Our data also supports previous findings describing the involvement of type I interferon response in the recruitment and activation of myeloid cells in the heart after MI ^14^, as we observed a massive recruitment of neutrophils with an IFN-response signature in the heart early after MI.

Consistent with previous reports^13,16^, cardiac-infiltrating neutrophils exhibited a profound tissue-specific signature, characterized by the local intensification of cardiac-specific markers and the acquisition of a SiglecF^+^ profile restricted to the myocardial environment. Intriguingly, while the SiglecF protein was exclusive to the heart, we observed that peripheral neutrophils—specifically those in the circulation and the spleen, but not the bone marrow—initiated the upregulation of the *Siglecf* transcript post-MI, suggesting a multi-step priming process that begins prior to extravasation. SiglecF^+^ neutrophils were found to be derived from GM-CSF- and TGFβ1-stimulated conventional neutrophils in chronic kidney disease associated with fibrosis^17^, and induced by TGFβ in pulmonary fibrosis^47^. In line with this observation, transcripts for receptors involved in SiglecF^+^ neutrophil differentiation *Tgfbr1*, *Tgfbr2*, and *Csf2ra*, were up-regulated over time post-MI in spleen and blood neutrophils, but not in the bone marrow, suggesting that SiglecF^+^ neutrophils could originate from spleen neutrophils. However, we cannot exclude that the SiglecF^+^ phenotype in neutrophils can be induced by other factors. For instance, in lung carcinoma, soluble RAGE released in the circulation has been shown to activate osteoblasts and fuel SiglecF^+^ neutrophil supply in the tumor ^19^.

In line with previous evidence^16,48^, targeting Ly6G did not appear as a reliable strategy to induce local neutrophil depletion. In addition to escape mechanisms to Ly6G targeting^36,48^, it was suggested that Ly6G, a glycosylphosphatidylinositol (GPI)-linked receptor, can play functional roles in neutrophil biology and should not be considered only as a surface marker. Indeed, the use of low doses of anti-Ly6G did not induce neutropenia but impaired neutrophil recruitment to sites of inflammation by abrogating neutrophil migration through impaired β2-intergrin surface expression^49^. In addition, mice with Ly6G-deficient neutrophils displayed delayed kinetics of pathogen-infected tissue entry, although neutrophil motion speed was conserved^50^. We had previously shown that targeting Ly6G in mice increased the proportion of SiglecF^+^ neutrophils in the infarcted heart^16^. Using a new Ly6G targeting strategy first described by Boivin et al.^36^, we further confirmed that SiglecF^+^ neutrophil proportions were increased 3 days post-MI after Ly6G targeting. Parallel to these findings, we observed an enrichment of mature *Retnlg^hi^*neutrophils within the heart following Ly6G intervention. The proximity of their gene expression profile to that of bone marrow neutrophils indicates a state of persistent emergency granulopoiesis, characterized by the mobilization and infiltration of bone marrow-proximal mature subsets into the cardiac tissue.

Anti-Ly6G treatment increased interstitial fibrosis in the MI border zone, consistent with previous observations^5^. We therefore propose that both enrichment in SiglecF^+^ neutrophils at early timepoints and accumulation of bone marrow-proximal mature-*Retnlg^hi^*neutrophils could contribute to increased fibrosis after Ly6G targeting. Interestingly, periodontitis was recently shown to increase cardiac fibrosis after MI by predisposing bone marrow neutrophils to convert into a SiglecF^+^ state, and by increasing their infiltration into the heart ^40^. SiglecF^+^ neutrophils were also associated with fibrosis in other pathological contexts, such as kidney fibrosis, where SiglecF^+^ neutrophils in mice, and Siglec-8^+^ neutrophil in humans, were detected in fibrotic kidney regions^17^. Moreover, SiglecF^+^ neutrophils were suggested to acquire the ability to produce collagen in kidney fibrosis and periodontitis contexts^17,40^. However, we did not find any evidence of direct collagen production by SiglecF^+^ neutrophils in our setting, suggesting a more indirect role in ECM remodeling. Neutrophils are well known for being actively involved in matrix remodeling by producing matrix metalloproteinases in cancer^51^ and MI^52^ contexts. Indeed, they presented the highest score for matrisome-related secretory factors among post-MI cardiac cells. Here, we show that SiglecF^+^ neutrophils can directly communicate with fibroblasts, especially through secretion of TGFβ and Cyclophilin A, to increase fibroblast wound healing capacities. Evidence that neutrophils serve as a significant source of CypA underscores the therapeutic importance of this axis in cardiac ischemic injury. Because CypA is a known driver of structural remodeling and hypertrophic growth, its neutrophil-specific secretion likely contributes to the deleterious outcomes observed in post-infarction heart failure^53^. Interestingly, elevated levels of Cyclophilin A were found in acute ST-segment elevation MI patients, associated to poor predictive outcome^54^.

Beyond direct neutrophil effects, Ly6G targeting reshaped the cardiac immune landscape, favoring the recruitment of monocytes and γδ T cells, presumably through enhanced chemotactic signaling. We identified γδ T cells as key mediators of this fibrotic response, as they significantly augmented fibroblast activation and proliferation. Given the known detrimental impact of γδ T cells on cardiac remodeling, their recruitment likely serves as a primary mechanism by which Ly6G-mediated neutrophil reprogramming accelerates pathological fibrosis ^41,55^. Moreover, intercellular communication analysis suggest Klk8/F2r ligand/receptor pathway involved in γδ T cells communication with fibroblasts. This pathway would be of interest as a target as PAR1 (encoded by *F2r*), highly expressed by fibroblasts, was previously shown to mediate post-MI remodeling response ^42,45,56,57^ and clinical trials targeting PAR1 using Vorapaxar showed interesting results in secondary prevention of atherothrombotic events^58^.

Ultimately, our data establishes that the orchestration of cardiac repair is fundamentally shaped by neutrophil heterogeneity. The observation that anti-Ly6G-mediated reprogramming promotes fibrosis via SiglecF^+^ subsets and γδ T cell recruitment redefines our understanding of the post-MI inflammatory landscape. Moving beyond broad-spectrum depletion, our findings suggest that the targeted redirection of neutrophil functional states represents a sophisticated therapeutic frontier for attenuating adverse remodeling and enhancing myocardial recovery.

## Acknowledgements

We thank the Single-Cell Center Würzburg and the histology and cytometry core unit facilities at PARCC.

## Conflict of interest

None.

## Financial support

This work was supported by the German Research Foundation (DFG SFB1525 project number 453989101, project to C.C. A.-E.S., A.Z., and seed grant program within the SFB1525 to M.P.), by Fondation pour la Recherche Medicale (FRM project number ARF202409019450 to M.P.), by the Université Paris Cité IdEx – inIdEx consortium CITY to M.P., by Agence Nationale de la Recherche (ANR-23-CPJ1-0134-01 to C.C.)

## Author contributions

Study conceptualization and design: M.P., C.C.; Data acquisition and analysis M.P., G.R. J.E.-K., M.Ga., M.Ge., L.T., C.C.; A.R., S.R.B., E.T.S., T.K., A.M.L.; Funding acquisition: M.P, A.Z., A.-E.S., C.C; Supervision: M.P., C.C.; Visualization: M.P., C.C.; Writing - original draft: M.P, C.C.; Review & editing: all authors

## Notes

### Competing Interest Statement

The authors have declared no competing interest.

