## Supplementary Figures for "Neutrophil terminal programming in the ischemic heart drives fibrosis after myocardial infarction"

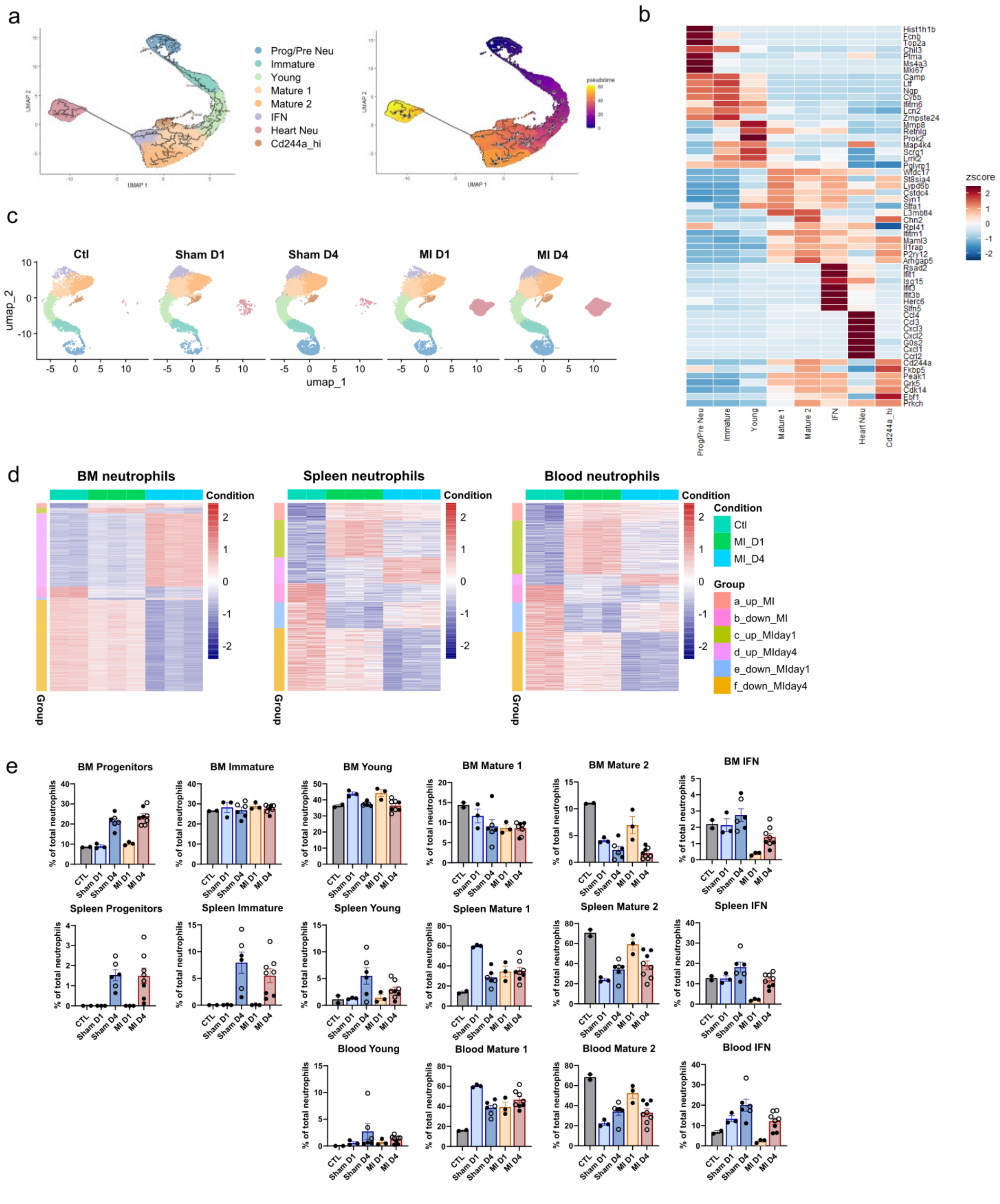

**Supplementary Figure 1: related to Figure 1. a)** UMAP plot generated by Monocle analysis (left panel) and pseudotime projected on neutrophils clusters (right panel). **b)** Top 7 markers genes for each cluster from Figure 1b calculated as z-score for variable features **c)** UMAP plot visualization of neutrophils according to experimental condition of origin. **d)** heatmap of pseudobulk-identified up- and downregulated genes in MI conditions compared to control in the bone marrow, spleen and blood. Ctl n= 2, MI D1 n=3, MI D4 n=3, male mice. **e)** neutrophil clusters proportion within total neutrophils in the blood, spleen and bone marrow according to experimental condition. One circle represents one mouse, open circles represent female mice. BM: bone marrow, MI: myocardial infarction.

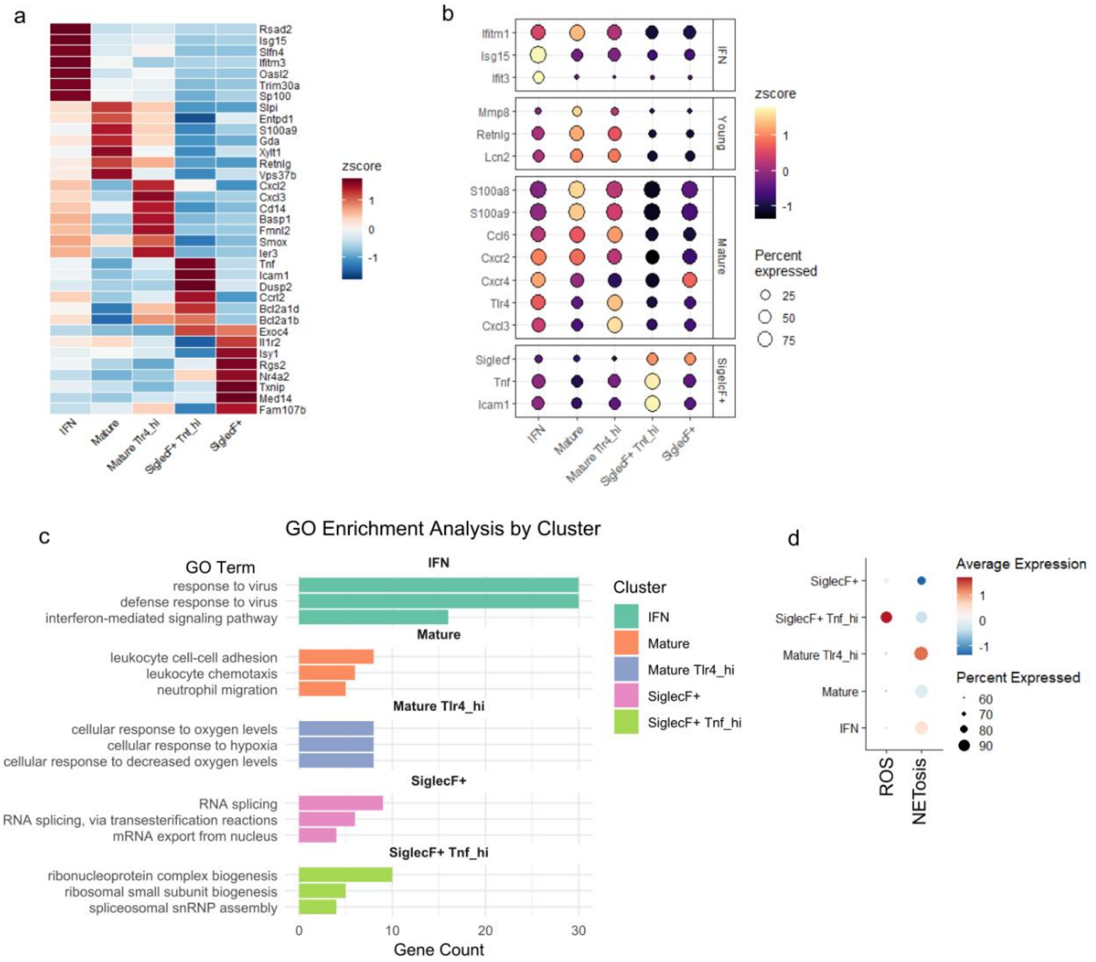


**Supplementary Figure 2: related to Figure 2. a)** Top 7 markers genes for each cluster from Figure 2a. **b)** Relative expression of the main marker genes used to identify neutrophil clusters in Figure 2a. **c)** Gene count for the top 3 gene ontology biological processes enrichment analysis in the indicated cardiac neutrophil cluster (all with adjusted p-value <0.05). **d)** ROS production and NETosis scores in indicated cardiac neutrophil clusters. GO: gene ontology, ROS: reactive oxygen species, NET: neutrophil extracellular trap.

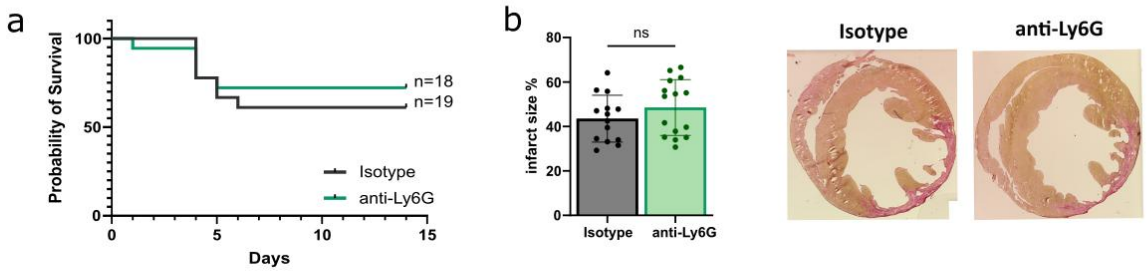


**Supplementary Figure 3: related to Figure 3. a)** Probability of survival over 14 days post-MI in the indicated experimental groups **b)** Infarct size, expressed as percentage of total left ventricular size, determined 14 days post-MI by picrosirius red staining; right panel: representative image for isotype and anti-Ly6G group, magnification 25X. Bar plots: one circle represents one mouse. Statistical tests: (panel b) unpaired t-test (ns=non-significant).

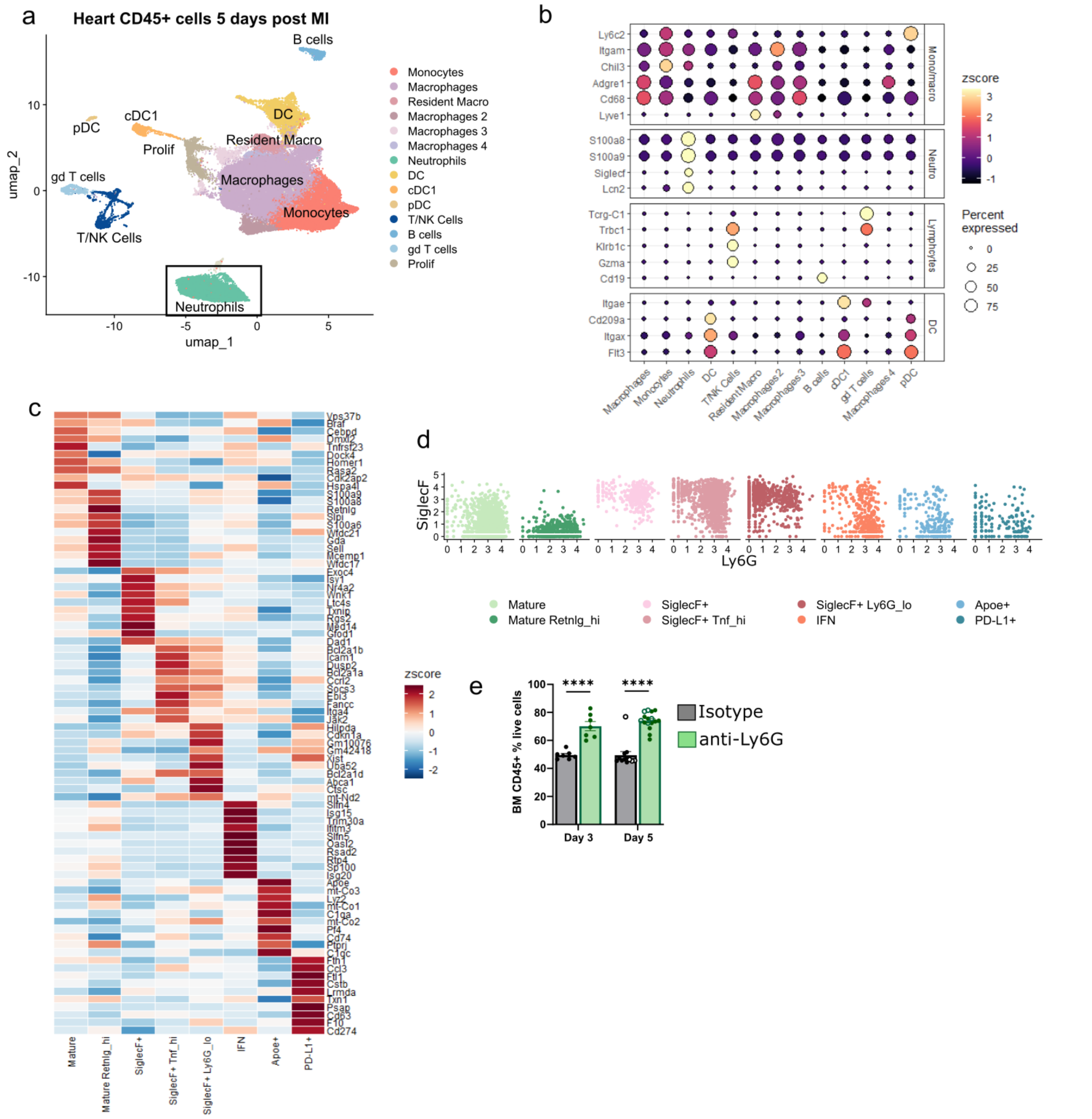


**Supplementary Figure 4: related to Figure 4. a)** UMAP visualization of total heart CD45$^+$ cells 5 days post-MI from the experimental setup detailed in Figure 3d. **b)** relative expression of the main marker genes used to identify CD45$^+$ cell clusters from panel a). **c)** Top 7 markers genes for each cluster from Figure 3e. **d)** CITE-seq signal for SiglecF vs. Ly6G in the indicated heart neutrophil populations identified in Figure 3e. **e)** CD45$^+$ cell percentage within total live cells in the bone marrow 3 and 5 days post-MI, evaluated by flow cytometry. Bar plots: one circle represents one mouse, open circle represent female mice. Statistical tests: (panel e) unpaired t-test (*<0.05; **<0.01; ***<0.001; ****<0.0001).

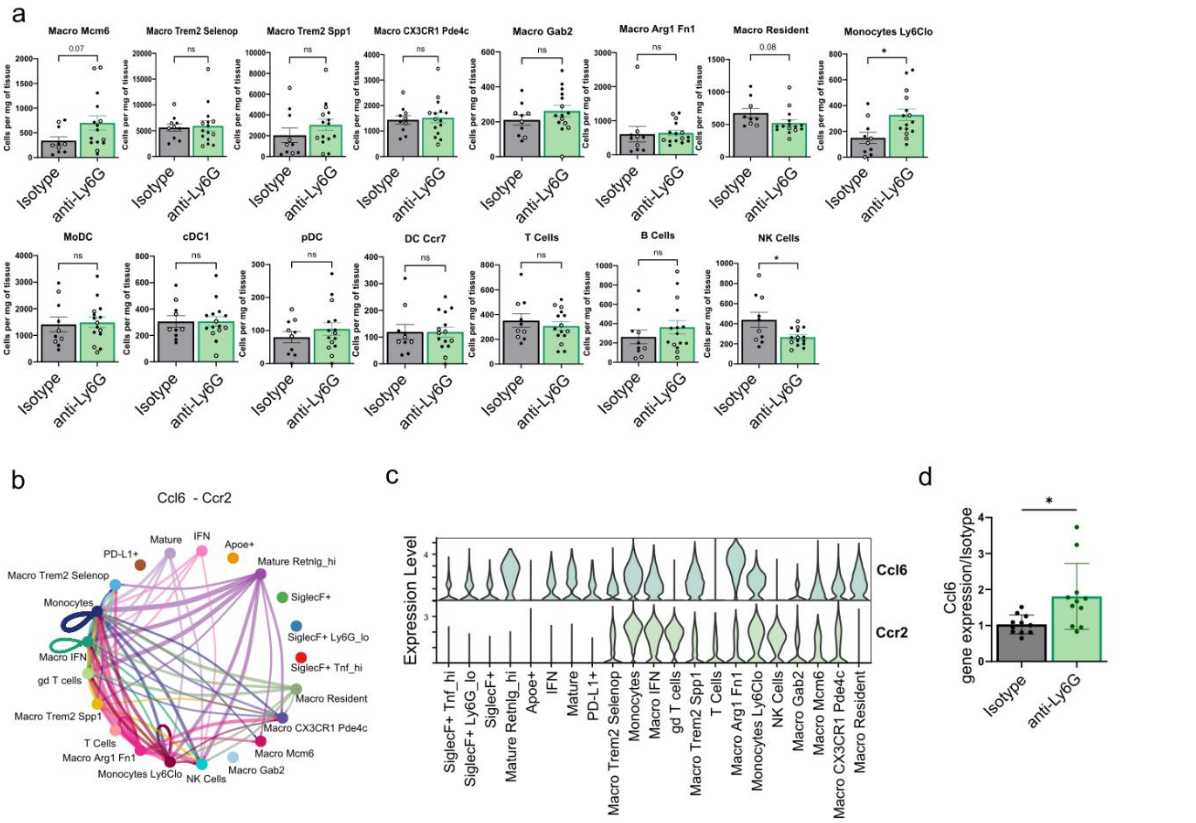


**Supplementary Figure 5: related to Figure 5. a)** Cell number per mg of heart tissue of the indicated cell clusters identified in Figure 5a. **b)** Probability of communication mediated by *Ccl6-Ccr2* pair between the indicated CD45+ heart cells from Figure 5a identified by CellChat analysis. **c)** *Ccl6* and *Ccr2* expression across the indicated CD45+ heart cells from Figure 5a. **d)** *Ccl6* expression assessed by quantitative PCR in heart tissue 3 days post-MI in isotype and anti-Ly6G groups. Bar plots: one circle represents one mouse, open circles female mice. Statistical tests: (panel a,d) unpaired t-test (*<0.05, **<0.01).

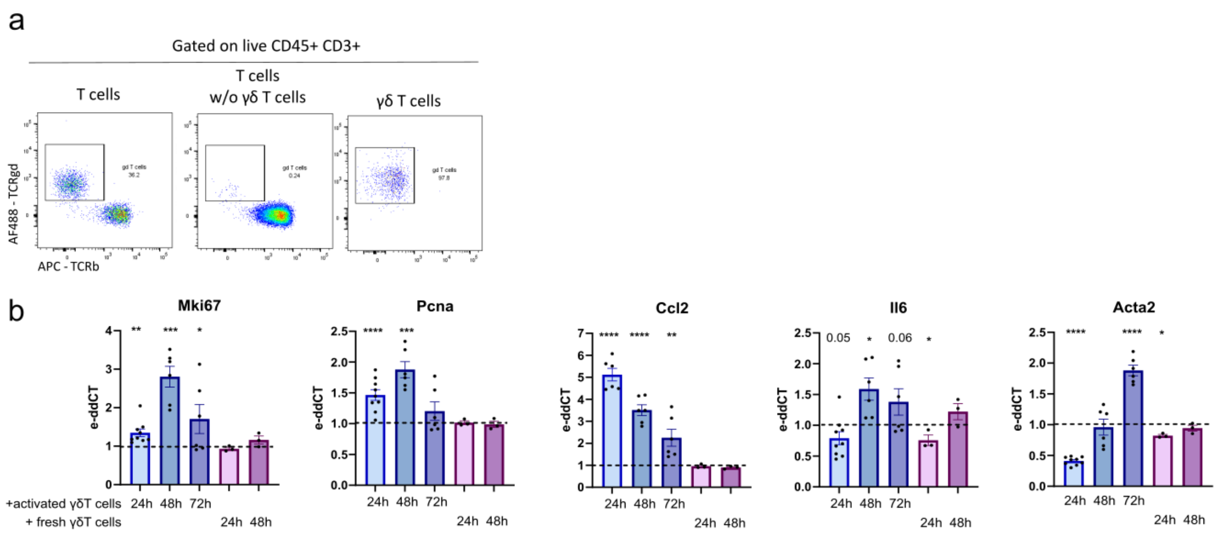


**Supplementary Figure 6: related to Figure 5. a)** γδ T cells purity after isolation **b)** Expression of the indicated genes by 3T3 fibroblasts co-cultured with activated or unstimulated γδ T cells for 24h, 48h and 72h. Gene expression was calculated by the ΔΔCt method, normalized to untreated time matched fibroblasts. Statistical test: one-way ANOVA (*<0.05; **<0.01; ***<0.001; ****<0.0001).
